# GAPMs form a heterotrimeric complex bridging the gliding machinery and the cytoskeleton across Plasmodium species

**DOI:** 10.64898/2026.03.27.714866

**Authors:** Akancha Mishra, Giedrė Ratkevičiūtė, Amy Ibrahim, Dušan Živković, Mohammad Zeeshan, Declan Brady, Andrew Bottrill, Jani R. Bolla, Eelco C. Tromer, Robert W. Moon, Rita Tewari, Clinton K. Lau

## Abstract

Apicomplexan parasites, such as malaria-causing *Plasmodium* spp., use a specialised actomyosin motor system known as the glideosome to drive movement through host tissue and invade host cells. This system is anchored to the inner membrane complex (IMC), a series of flattened vesicles located beneath the plasma membrane, and thought to be linked to the underlying cytoskeleton by the GAPM protein family. However, it is not known how these GAPM proteins are localised across the *Plasmodium* life cycle, and whether different family members function alone or together. Here, we show that in two *Plasmodium* species GAPM2 is an IMC component whose recruitment and organisation are tightly coordinated with nuclear and cytoskeletal dynamics during parasite replication and differentiation. We find that the GAPM2 interactome remodels between asexual and sexual stages using mass spectrometry. To understand the molecular relationship between three GAPM paralogues, we solved a cryo-electron microscopy structure of the GAPM complex. This revealed an obligate heterotrimeric architecture that forms an asymmetric platform, likely to serve as a docking interface for other components of the glideosome. Finally integrating our GAPM heterotrimer structure with mass spectrometry data allowed us to propose a unified structural model of the glideosome that is conserved across apicomplexan parasites.

## Introduction

Despite global control efforts, malaria-causing *Plasmodium* protozoan parasites continue to pose a major public health challenge, resulting in 610,000 deaths in 2024^1^. These obligate parasites, and other apicomplexans, have evolved specialised machinery to survive within their hosts. One distinctive example is the glideosome, an actomyosin-based multiprotein complex that drives gliding motility through host tissue and cell invasion. The glideosome is contained within the parasite pellicle that comprises the plasma membrane (PM) and the inner membrane complex (IMC), a series of cortical flattened vesicles beneath the PM, which together with microtubules form the cytoskeleton^2,3^.

The motor complex within the glideosome is anchored to the PM-facing IMC surface. It comprises an atypical class XIV myosin, MyoA, which lacks a canonical tail domain. MyoA binds two essential light chains, MTIP (known as MLC1 in *P. berghei)* and ELC^4–9^. MTIP interacts with glideosome-associated protein (GAP) GAP45^10^, which in turn binds GAP40 and GAP50^11^.

The link between glideosome and cytoskeleton is less well-characterised. Studies have implicated glideosome-associated proteins with multiple membrane spans (GAPMs), a family of integral membrane proteins that are conserved across apicomplexan parasites ^12–14^. *Plasmodium* species encode three GAPM paralogues, GAPM1, GAPM2, and GAPM3, which have previously been shown to coprecipitate (Rayavara et al., 2009) and have been suggested to form large oligomers^12^. GAPMs reside within the inner membrane of the IMC, with their N- and C- termini facing the cytoplasm (Harding et al., 2019). Several proteomic analyses suggest that GAPMs form part of the scaffold bridging the glideosome to the cytoskeleton^5,12,15–17^.

Functional studies support this structural role for GAPMs: in *Toxoplasma gondii*, conditional degradation of either GAPM1a or GAPM2a disrupts both IMC morphology and cortical microtubule organisation^13^, while knockdown of GAPM2 in *P. falciparum* results in a severe invasion defect^17^. In *P. berghei*, GAPM genes appear to be transcribed together at high levels in the asexual intraerythrocytic stage and during sexual development through the gametocyte and ookinete stages ^18^. However, how this translates to protein levels is unclear: How do GAPMs coordinate with other aspects of cell development, and how do they fit into the glideosome? Do GAPM paralogues function alone or together?

In this study, we examine the relationship between GAPMs, the glideosome and the microtubule network in two different *Plasmodium* parasites and across different stages of the life cycle using expansion microscopy and mass spectrometry. We show that GAPMs assemble into a defined, stable heterotrimer *in vitro* using a combination of native mass spectrometry and cryo-EM. The residues involved in the inter-subunit contacts are conserved, with the arising asymmetry presenting interfaces that will bind other glideosome and IMC components. Together, our data suggest a model for the organisation of heterotrimeric GAPM complex within the IMC and support their role in anchoring the glideosome in *Plasmodium* parasites.

## Results

### GAPM2 association with the IMC during nuclear division and cell differentiation

Initially we examined the expression and location of GAPM2 during development at different stages of the *Plasmodium* life cycle. To do this we generated a GAPM2 C-terminal GFP fusion (Extended Data Fig. 1A-C), to monitor its localisation in *Plasmodium berghei* (*Pb*) using live-cell microscopy (Fig. 1). In the asexual blood stage, GAPM2 was detectable from the late ring stage as a small cytoplasmic vesicle and increased in abundance in late trophozoites, where multiple cytoplasmic vesicles were observed, indicating synthesis prior to nuclear division (Fig. 1A, Extended Data Fig. 1D). At the onset of schizogony, GAPM2 was visible as plaques at the parasite periphery associated with individual nuclei, and in mature segmented schizonts each merozoite was surrounded by a GAPM2-enriched IMC (Fig. 1A).

**Figure 1.**
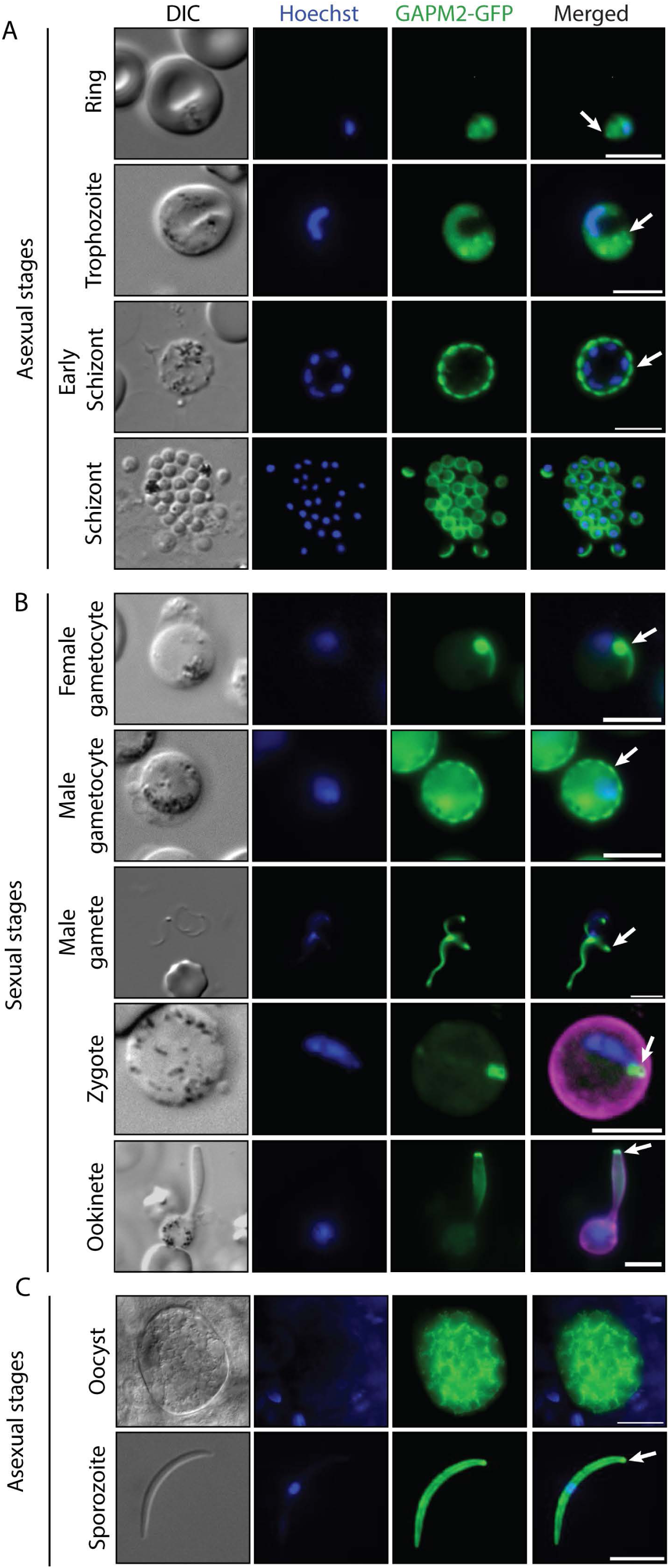
Dynamic localization of GAPM2 throughout the *Plasmodium berghei* life cycle. Representative live-cell fluorescence images of parasites expressing a C-terminal GAPM2-GFP fusion across asexual and sexual developmental stages. **(A)** In asexual stages, *Pb*GAPM2-GFP appears as puncta in rings and trophozoites (arrows), redistributing to the periphery to outline the IMC in schizonts (arrow). **(B)** In sexual stages, *Pb*GAPM2-GFP localizes to the surface of activated male gametocytes (arrow) and adopts an apical ring-like pattern in female gametocytes, zygotes, and in elongating ookinetes (arrow). **(C)** *Pb*GAPM2-GFP is also observed in sporulating oocysts (scale bar 20 µm) and localizes to the apical tip (arrow) and along the IMC of salivary gland sporozoites. Scale bars 5 µm.

During sexual development, GAPM2 had distinct localisations in both activated male and female gametocytes, forming a discontinuous surface localisation in males and a punctum in females. It persisted in activated male gametes despite their reliance on a flagellum for movement, as opposed to the glideosome (Fig. 1B). Following fertilisation, in zygotes, GAPM2 was present in a location similar to that in the female gametocyte, but during differentiation into ookinetes the signal became mainly restricted to the apical end of the growing protuberance. GAPM2 was also observed in sporulating oocysts and around the periphery of salivary gland sporozoites (Fig. 1C). Overall, these data indicate that GAPM2 is present during major stages of the parasite life cycle.

To understand better the spatiotemporal dynamics of GAPM2 during asexual blood stage cell division, we imaged schizogony at several timepoints (Fig. 2). *Pb*GAPM2-GFP first appeared at the parasite periphery at the onset of nuclear division and remained associated with the expanding IMC membranes as daughter merozoites begin to differentiate (Fig. 2B). To determine whether these dynamics are conserved across species, we generated a *P. knowlesi* (*Pk*) GAPM2-mNG parasite line (Extended Data Fig. 2), which showed a similar distribution as that of *Pb*GAPM2 during schizogony (Fig. 2C). Super-resolution structured illumination microscopy (SIM) confirmed continuous membrane association of GAPM2 throughout *P. berghei* schizont maturation (Fig. 2D).

**Figure 2.**
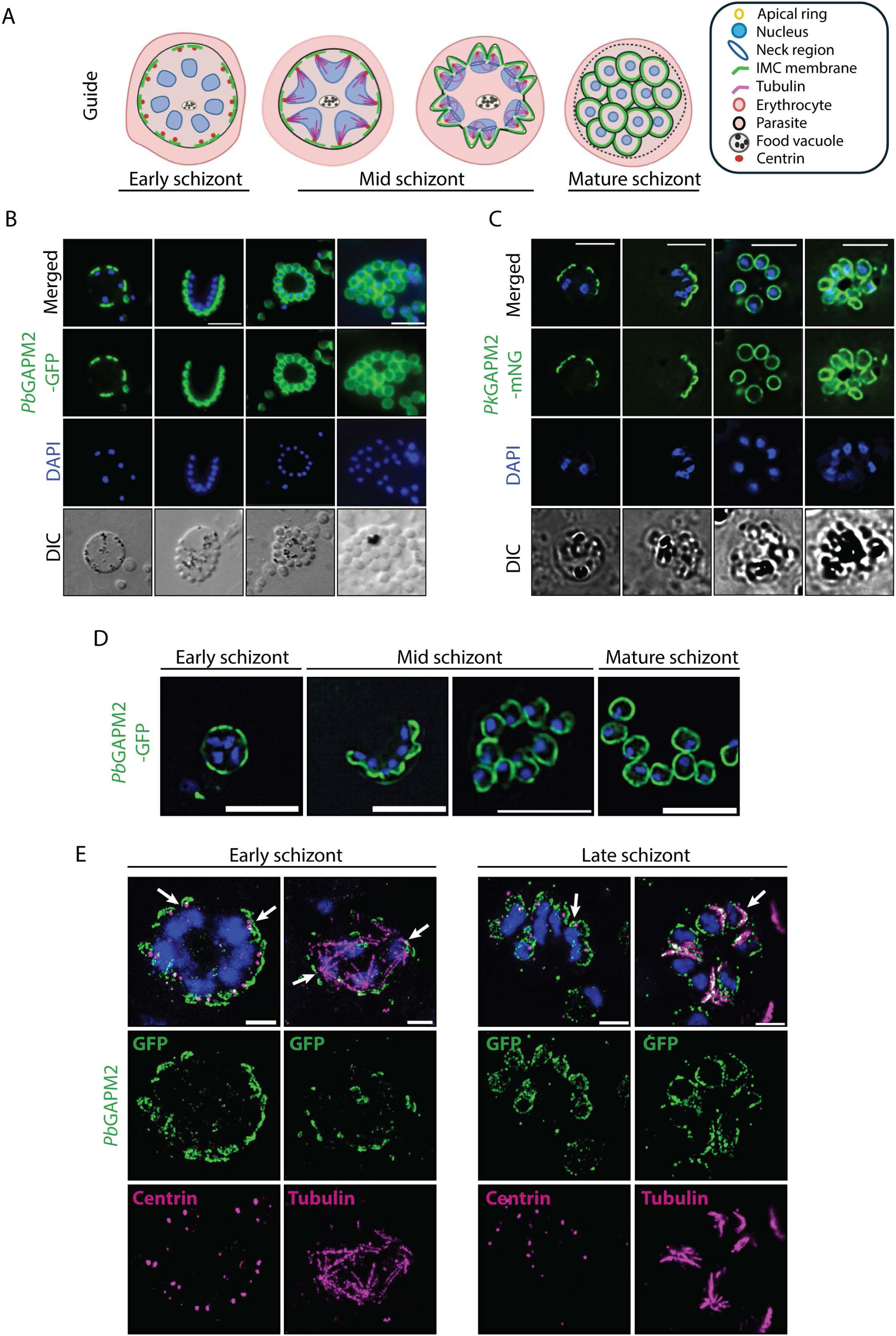
GAPM2 marks IMC assembly and coordinates with nuclear and cytoskeletal elements during schizogony. **(A)** A schematic for the asexual development of blood stage parasites tracking the GAPM2 expression (green) along with cytoskeleton markers like centrin (red) and tubulin (magenta) and the neck region (blue). **(B)** For the live-cell imaging of GAPM2-GFP in *P. berghei* shows early surface localization at the onset of nuclear division, with GAPM2-enriched IMC membranes elongating as nuclei segregate, culminating in mature schizonts in which each nucleus is enclosed by GAPM2-positive IMC. **(C)** A similar pattern is observed in *P. knowlesi* schizonts, with early surface enrichment expanding to cover the parasite in mature stages. **(D)** SIM confirms *Pb*GAPM2 localization along IMC membranes throughout schizogony. Scale bars 5 µm. **(E)** U-ExM of *Pb*GAPM2 GFP co-stained with centrin-GFP (magenta-green) or tubulin-GFP (magenta-green) in early and mature schizonts reveals the coordinated organization of nuclear, centromeric, and cytoskeletal elements during schizont development in *Plasmodium* asexual blood stages (arrows). All scale bars 5 µm.

We examined the interplay between GAPM2, cytoskeleton and nuclear dynamics using ultrastructural expansion microscopy (U-ExM). Early and mature *Pb*GAPM2-GFP schizonts were co-stained for centrin (an outer component of the bipartite microtubule organising centre [MTOC]) and tubulin (cortical microtubules and mitotic spindle). From the onset of early schizogony, each dividing nucleus was associated with two newly formed GAPM2-positive IMC structures positioned adjacent to duplicated MTOCs (Fig. 2A,E). Microtubules emanating from the MTOC likely facilitate nuclear segregation, indicating that IMC biogenesis is coordinated with MTOC duplication and cytoskeletal assembly prior to completion of nuclear division. As schizogony proceeded, each nucleus segregates together with its associated IMC, which expanded down the length of each nascent merozoite (Fig. 2A,E). Collectively, these data from different parasite stages and species show that GAPM2 is a conserved IMC component whose recruitment and organisation are tightly coordinated with nuclear and cytoskeletal dynamics during parasite replication and differentiation.

### *Pb*GAPM2 spatiotemporal dynamics during gametocyte activation and ookinete development

Next we focused on GAPM2 localisation during sexual development. We purified *Pb*GAPM2-GFP gametocytes and monitored the protein by live-cell imaging. GAPM2 was not detected in non-activated gametocytes, but upon activation GAPM2 rapidly appeared at the periphery of both male and female gametocytes (Fig. 3A,B). Subsequently cells were fixed at 4-8 min post-activation (pa) and analysed by SIM. In male gametocytes, IMC vesicles appeared fragmented, with apparent gaps between vesicles. Interestingly, we observed a variable number of IMC vesicles (7-9) per differentiating MTOC (8^19^)at 8 min pa (Fig. 3C), in contrast to the strict two IMC vesicle per MTOC ratio in asexual stages (Fig. 2B, E). Each male gamete retained GAPM2 signal along its length, with enrichment at its tip (Supplementary Video 1). In female gametocytes, GAPM2 localisation progressed from a clearly-defined arc into a complete ring (Fig. 3C).

**Figure 3.**
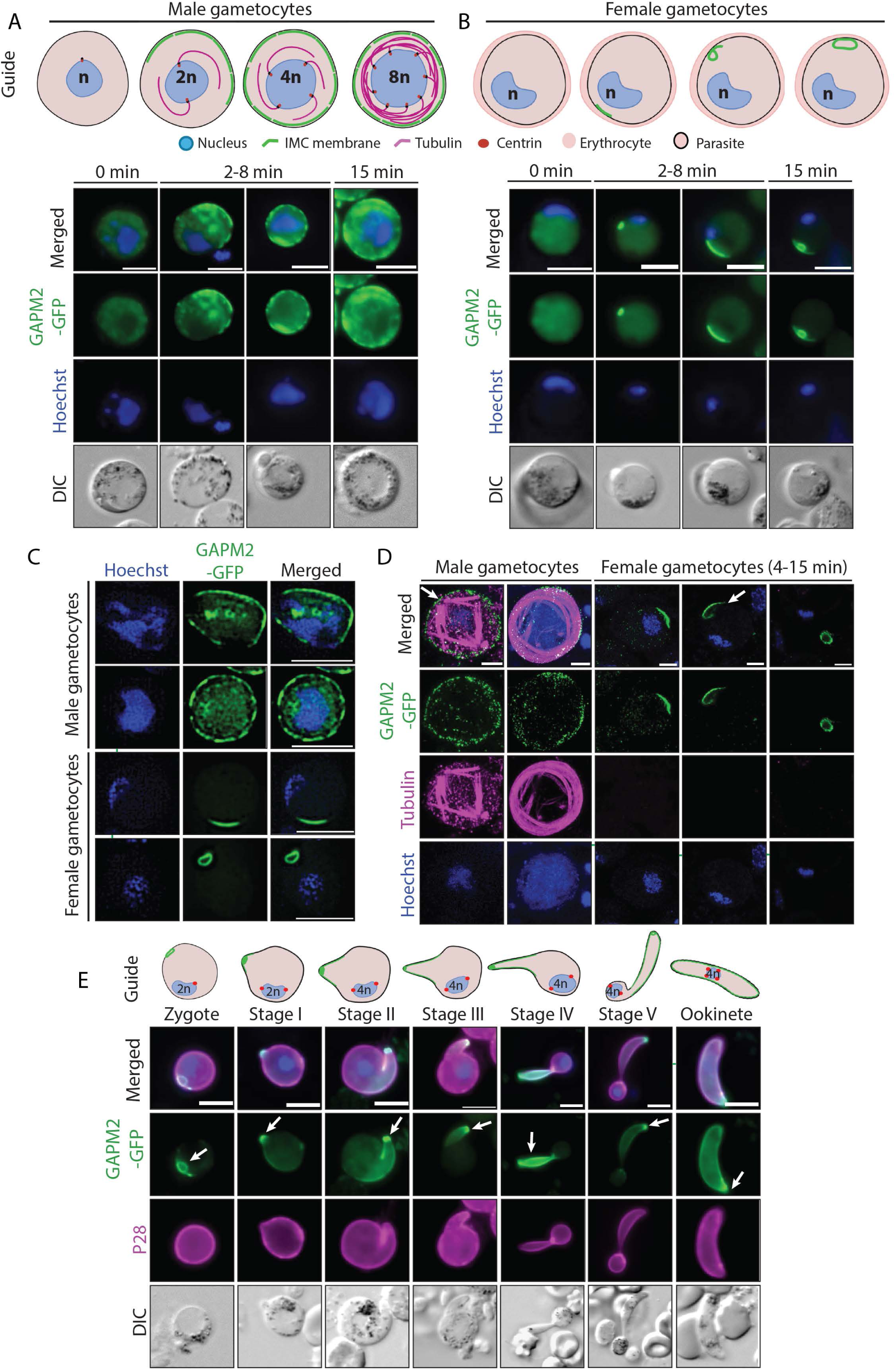
GAPM2 marks IMC assembly and apical polarity during sexual development and ookinete differentiation. **(A)** Live-cell imaging of purified gametocytes shows that *Pb*GAPM2 is absent in non-activated male and female gametocytes but rapidly appears at the surface of both sexes upon activation. **(B)** Super-resolution SIM of 4-8 min post-activation gametocytes reveal fragmented IMC vesicles in males while females exhibit *Pb*GAPM2 as a forming apical arc that develops into a ring. **(C)** U-ExM with tubulin (magenta) and GFP (green) staining demonstrates alignment of *Pb*GAPM2-positive IMC structures on the surface in males (arrows) and establishment of apical polarity in females (arrows). **(D)** After fertilization, *Pb*GAPM2 remained concentrated at the apical polar ring (arrows), which elongated during parasite protrusion and ultimately extended to form the crescent-shaped ookinete. At this stage, *Pb*GAPM2 was distributed across the entire IMC membrane and densely accumulated at the apical end. All scale bars 5 µm.

We stained gametocytes at 4-8 minutes pa for tubulin and GFP, using U-ExM to examine the coordination of IMC assembly and the tubulin-based cytoskeleton. In males, tubulin arrays extended underneath the fragmented GAPM2-positive IMC structures, whereas in females, the location of GAPM2 marked the establishment of subsequent apical polarity (Fig. 3D). Expanded gels were additionally stained with NHS-ester to assess the spatial relationship between GAPM2-enriched IMC, basal bodies and MTOCs (Extended Data Fig. 1E). However, no clear spatial correlation between these structures could be resolved. In female gametocytes, NHS-ester staining revealed no features that could explain the initiation of polarity in unfertilised cells (Fig. 3D).

The GAPM2-labelled ring observed in the female gametocyte was retained in the fertilised zygote and persisted throughout *ex vivo* ookinete differentiation. GAPM2 remained enriched at the apical end of the developing ookinete protrusion as the IMC extended down the length of the elongating retort, resulting in a mature ookinete encapsulated in a GAPM2-positive IMC (Fig. 3E). This suggests that GAPM2 linked to apical specification, similar to during schizont development. However, the fact that GAPM2 in gametocytes appears less coordinated with cytoskeletal dynamics from gametocytes led us to hypothesise that its interactome may change in different life cycle stages.

### Differential Interactions of GAPM2 define a conserved IMC-glideosome module

Whilst previous studies have placed GAPM2 in the glideosome, we wanted to obtain a wider view of the GAPM2 interactome and investigate how it might vary in different life cycle stages. To do this we performed pull-down mass spectrometry analysis using GAPM2 as a bait, including crosslinker prior to lysis to enrich for distal or transient interactors. For *Pb*GAPM2-GFP, parasite lysates were prepared from schizont and gametocyte stages, whereas for *Pk*GAPM2-mNG parasites were tightly synchronised to isolate late-stage schizonts.

Across all datasets, GAPM2 associated with other members of the glideosome, as well as IMC proteins including PhIL1 and its interacting partners (Fig. 4A-C). The magnitude of ΔriBAQ values across datasets is consistent with a proximity-based interaction hierarchy centred on GAPM2. Applying a stringent threshold, schizont-stage data from *P. berghei* showed strong associations between GAPM2 and GAPM1, GAPM3, PhIL1, glideosome components GAP50, GAP40, MyoA and MTIP, with a lower relative abundance of GAP45 (Fig. 4A). We found a similar interaction profile in *P. knowlesi* schizonts, though GAPM1 and GAP45 are not consistently observed (Fig. 4B, Supplementary Data 2).

**Figure 4:**
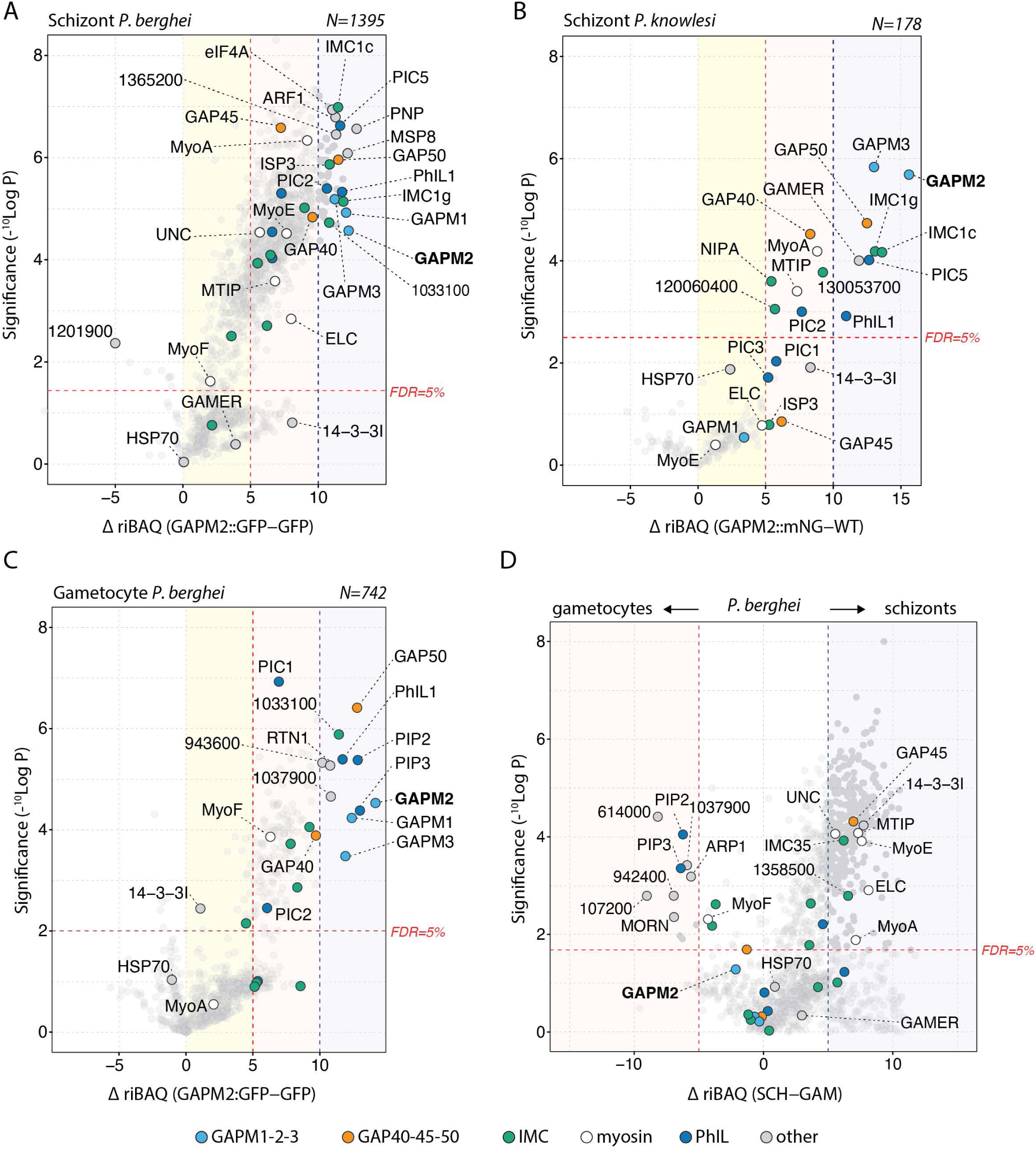
Stage-specific GAPM2 interactomes reveal conserved IMC-glideosome associations. **(A-C)** Volcano plots of protein abundance in GAPM2 pulldowns relative to controls across species and life cycle stages: **(A)** *P. berghei* schizonts (*Pb*GAPM2-GFP vs WT-GFP), **(B)** *P. knowlesi* schizonts (*Pk*GAPM2-mNG vs parental), and **(C)** *P. berghei* gametocytes at 1.5-6 min post-activation (*Pb*GAPM2-GFP vs WT-GFP) (n = 3). **(D)** Comparison of GAPM2-associated proteins between schizont and gametocyte stages in *P. berghei*, highlighting stage-dependent differences in enrichment. Shaded regions indicate ΔriBAQ intervals (light yellow, 0-5; red, 5-10; blue, >10). Higher ΔriBAQ values indicate increased relative abundance in GAPM2 pulldowns, consistent with proximity within the crosslinked complex and/or greater stability. Statistical significance was assessed using two-tailed unpaired Student’s t-tests with Benjamini-Hochberg correction (false discovery rate 5%; horizontal dashed line).

Comparing the *P. berghei* gametocyte data with the schizont stage datasets revealed pronounced stage-specific differences (Fig. 4C,D). Core glideosome components GAPM1 and GAPM3, GAP40 and GAP50 showed comparable relative abundance between stages, as are the proteins attributed to the inner IMC membrane, such as IMC1g, IMC1c and ISC3, PhIL1 and some of its interactors such as PIC1. In contrast, GAP45 and myosin motor components, MyoA, MTIP and ELC are much reduced in the gametocyte samples. Additionally, PhIl1 interacting proteins PIP2 and PIP3 are increased compared with asexual stages. Together, these data define a GAPM2-associated core module and reveal stage-dependent remodelling of motor-associated components during early gametocytogenesis.

### GAPMs assemble into a defined 1:1:1 heterotrimer *in vitro*

Given the co-enrichment of GAPM1 and GAPM3 in GAPM2 mass spectrometry data, we investigated whether the three GAPM paralogues interact directly with one another. AlphaFold predicted a trimeric architecture but assigned similar confidence to homotrimeric (iPTM_TM_ 0.83-0.92) and heterotrimeric (iPTM_TM_ 0.92) models (Extended Data Fig. 3). Therefore, to test predictions for GAPM assembly, we recombinantly expressed GAPM proteins.

Expression of individual *P. falciparum* GAPMs gave low yields, protein aggregation, and poorly-resolved size-exclusion chromatography (SEC) profiles (Fig. 5A), suggesting instability in isolation. In contrast, co-expression of the three GAPMs yielded a single, monodisperse species containing all GAPMs (Fig. 5A), suggesting assembly of a stable complex. To further understand the subunit stoichiometry, we subjected the sample to native mass spectrometry analysis. The resultant spectrum showed a major charge-state series corresponding to the measured mass of 115,346 ±8 Da, which is in excellent agreement with the theoretical mass of a 1:1:1 GAPM1/GAPM2/GAPM3 complex (35,033; 46,413; and 33,882 kDa, respectively), including affinity tags (Fig. 5B). Additional adduct peaks seen were consistent with co-purified lipids, but no alternative assemblies corresponding to homo- or hetero-oligomers, or monomers were detected. Together, these results demonstrate that GAPM1, GAPM2, and GAPM3 assemble into a defined 1:1:1 heterotrimeric complex, providing a molecular explanation for the co-occurrence of GAPM paralogues in our interactome analyses.

**Figure 5.**
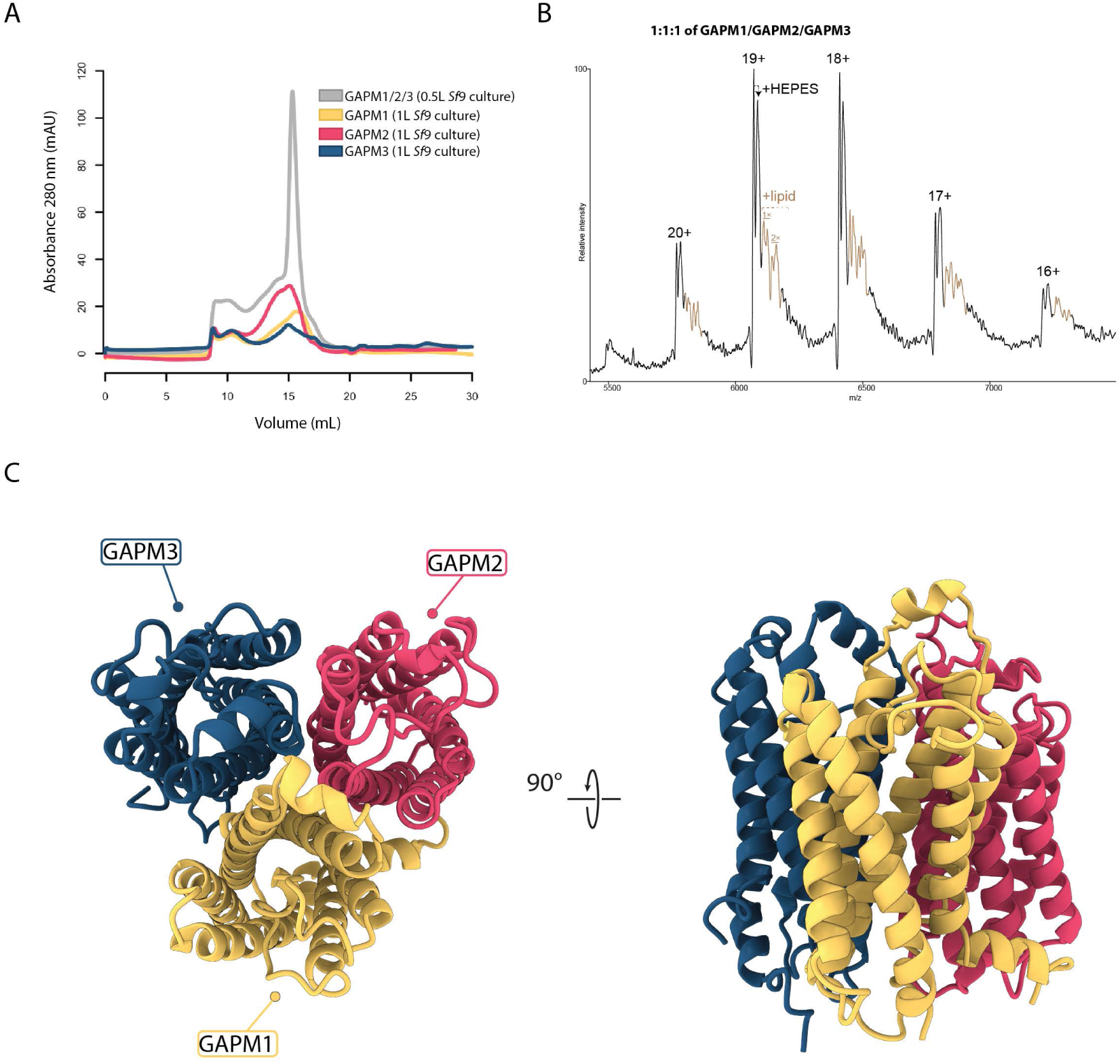
*Pf*GAPM1, *Pf*GAPM2, and *Pf*GAPM3 form an obligate heterotrimer. **(A)** Size-exclusion chromatography (Superose 6 10/300) profiles of individually expressed *Pf*GAPM proteins (GAPM1, yellow; GAPM2, dark pink; GAPM3, dark blue) compared with the co-expressed GAPM1/GAPM2/GAPM3 complex (grey). Individual GAPMs were purified from 1 L *Sf*9 cultures, whereas the GAPM1/GAPM2/GAPM3 complex was purified from 0.5 L *Sf*9 culture. **(B)** Native mass spectrum of the GAPM complex collected from the monodisperse SEC peak in (A). A dominant charge-state series is observed whose measured mass is consistent with a 1:1:1 GAPM1/GAPM2/GAPM3 assembly. Additional higher-mass adduct peaks indicate co-purified/bound phospholipids (olive brown). Peak splitting in the main series and adducts is attributable to HEPES adduction carried over from purification buffers. **(C)** Cryo-EM reveals the architecture of the *Pf*GAPM heterotrimer. Individual chains are coloured according to protein identity: GAPM1 (yellow), GAPM2 (dark pink), and GAPM3 (blue).

### Cryo-EM structure reveals asymmetric architecture of the GAPM heterotrimer

To gain structural insight into the GAPM assembly, we determined the cryo-EM structure of the complex in dodecylmaltoside (DDM) detergent at an overall resolution of 3.3 Å (Fig. 5C, Extended Data Fig. 4, 6A, Supplementary Table 3). The density was well-resolved in the transmembrane region and in the loops that would face the IMC lumen (Extended Data Fig. 6). As in our native mass spectrometry data, there was evidence of copurifying lipids (Extended Data Fig. 4B). This map allowed unambiguous model building and assignment of each subunit (Extended Data Fig. 6C). We also solved a second structure in glyco-diosgenin (GDN) detergent (Extended Data Fig. 5, 6B), which closely resembled our primary map (Supplementary Table 3).

The structure reveals that GAPM1, GAPM2 and GAPM3 organise into a well-defined heterotrimer, each contributing a six transmembrane-helix (6TM) bundle (Fig. 5C). Superposition of the three subunits shows that their transmembrane cores are structurally similar (RMSD_TM_= 0.9-1.0 Å, Fig. 6B). However, examining the interfaces between the subunits reveals the basis for heterotrimer formation. The assembly is specified by a unique set of hydrogen bonds and salt bridges between GAPM subunits at the luminal and cytoplasmic interfaces (Fig. 6A). These interactions would not arise in homotrimers of any single GAPM, suggesting that the heterotrimeric arrangement arises from cognate interfaces between GAPMs in the transmembrane region.

**Figure 6.**
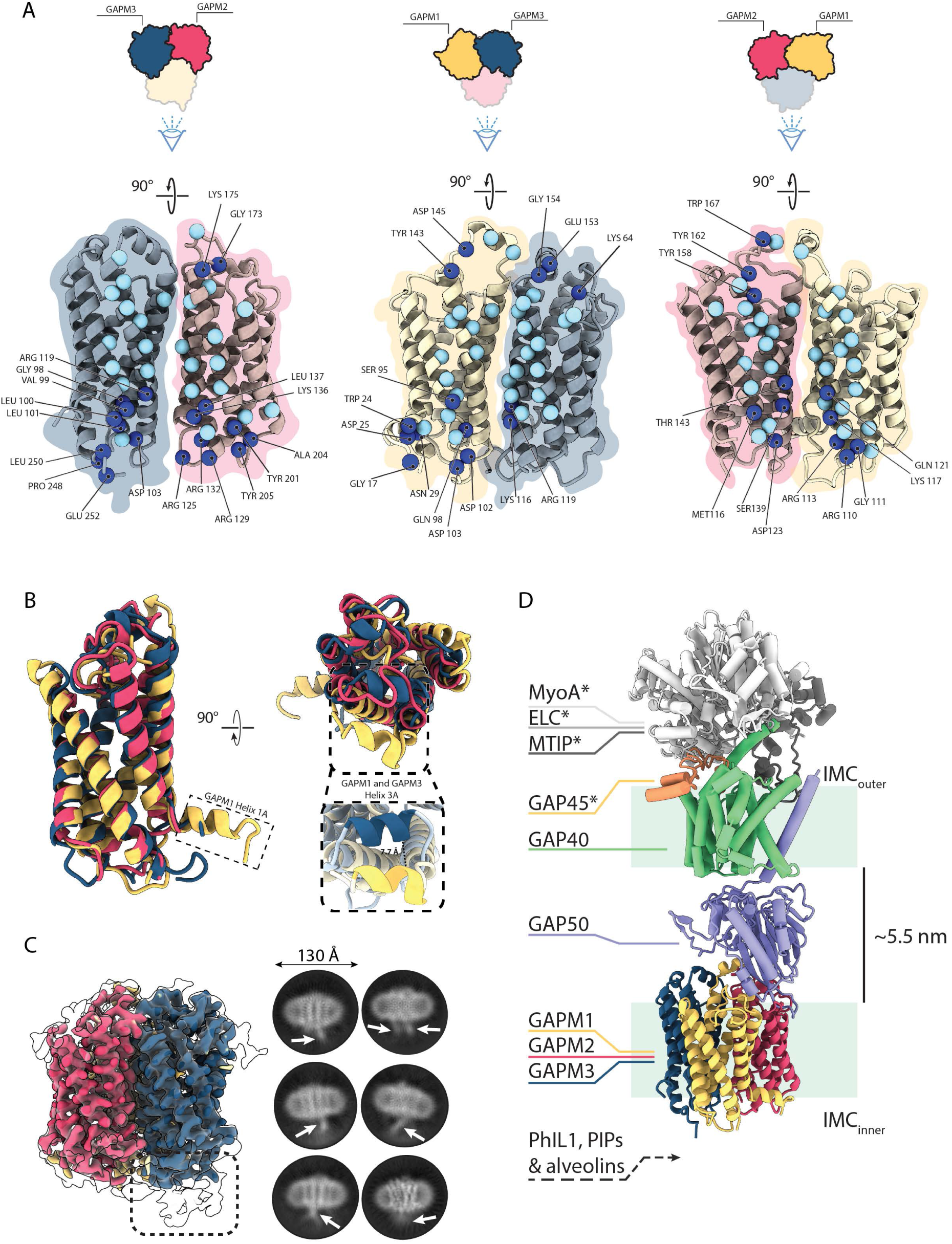
Asymmetric architecture, interaction interfaces, and flexible termini of the *Pf*GAPM heterotrimer. **(A)** Interaction interfaces within the GAPM heterotrimer, shown for the pairs GAPM1-GAPM2 interacting with GAPM3, GAPM1-GAPM3 interacting with GAPM2, and GAPM2-GAPM3 interacting with GAPM1. Dark blue spheres denote Cα positions of residues involved in hydrogen bonds and salt bridges with the adjacent subunit, while light blue spheres indicate hydrophobic contact sites. Cartoon diagrams of the GAPM1/2/3 top view are shown to aid clarity. **(B)** Superposition of GAPM1, GAPM2, and GAPM3 reveals high structural similarity. The RMSD across 182 pruned atom pairs is 1.006 Å (RMSD across all 221 pairs: 2.051 Å). The largest deviation occurs in a luminal face, with the GAPM1 helix 3A (residues 147-153) positioned 8.7 Å from the corresponding GAPM3 helix (residues 145-152). GAPM1 also contains an N-terminal amphipathic helix that is not observed in GAPM2 or GAPM3, termed GAPM1 helix 1A. **(C)** Cryo-EM density of the GAPM heterotrimer embedded in a DDM micelle. The coloured map shows the consensus reconstruction (0.175σ), while the overlaid translucent grey map at lower σ (0.0867σ) reveals additional density beneath the GAPM3 monomer. Representative 2D class averages are shown on the right; white arrows mark diffuse density that likely corresponds to the flexible N- and C-termini. **(D)** Composite AlphaFold model. The heterotrimeric GAPM cryo-EM structure is shown in cartoon, the AlphaFold2 predictions are shown in rod representation. Components marked with * are more enriched in the asexual blood stage compared to the gametocyte stage

To understand the conservation of these interfaces we interrogated orthologues of GAPM proteins in the PFAM database^20^. Here, proteins are found in many apicomplexans and the related chromerids, and are predicted to adopt a similar structure by AlphaFold (Extended Data Fig. 7). In apicomplexans, GAPM proteins partition into GAPM1-3 subfamilies, whereas chromerid GAPMs classify separately (Extended Data Fig. 7, Supplementary Data 4). Within each subfamily, residues which form side-chain hydrophilic contacts are conserved (Extended Data Fig. 8A, Supplementary Data 5,6,7). In agreement with this, co-expressing GAPM transmembrane constructs lacking their flexible N- and C- termini results in a stable heterotrimeric complex (Extended Data Fig. 9), showing that the intra-trimer interactions specify the heterotrimeric complex.

The distinct loops of each GAPM create asymmetric surfaces on the luminal and cytoplasmic faces. On the luminal face, the GAPM loops diverge significantly (Fig. 6B). Notably, GAPM1 and GAPM3 each contain an additional well-defined helix, referred to here as helix 3A (Fig. 6B, Extended Data Fig. 9C). In GAPM1 helix 3A extends further toward the central subunit interface than that of GAPM3, indicating differences in their luminal architectures. In contrast, GAPM2 lacks this helix, with cryo-EM density here suggesting an ordered loop. Within this asymmetric surface, GAPM2 and part of GAPM1 are better conserved amongst Apicomplexa (Extended Data Fig. 8B), suggesting this area is important for function.

The asymmetric cytoplasmic face is more conserved compared to the luminal face (Extended Data Fig. 8B). A striking feature is the presence of an amphipathic helix in GAPM1 (helix 1A), which appears to be absent in GAPM2 and GAPM3 (Fig. 6B). Additionally, diffuse density extending beyond the cytoplasmic face is visible in 2D class averages (Fig. 6C) and beneath GAPM3 at lower thresholds in our 3D reconstruction. We attribute this density to unmodelled N- and C-terminal extensions, suggesting these termini are dynamic.

### Integrating mass spectrometry and AlphaFold2 to position the GAPM heterotrimer in the glideosome

To investigate how an asymmetric, heterotrimeric GAPM complex would fit into the glideosome, we integrated our cryo-EM structure and mass spectrometry data to model the glideosome across the two membranes of the IMC using AlphaFold. We initially provided AlphaFold with all identified glideosome proteins, though this produced a poor model with the two transmembrane elements, the GAPM heterotrimer and GAP40/GAP50, adjacent to one another (Extended Data Fig. 10A).

As an alternative approach, we leveraged previous evidence^6–9,11,12^ to predict the glideosome in three overlapping segments (Extended Data Fig. 10B-D). Aligning these models produced an overall architecture that would span two membranes as anticipated, predicting an intermembrane distance of approximately 5.5 nm (Fig. 6D).

This model predicts that GAP50 is recruited to the GAPM heterotrimer through the luminal conserved patch formed by GAPM1 and GAPM2. GAP50 is anchored into the outer IMC membrane to GAP40, which in turn recruits GAP45 and the motor complex in asexual stages.

## Discussion

### GAPM heterotrimer is likely maintained across the *Plasmodium* lifecycle

Apicomplexa contain multiple GAPM proteins that are hypothesised to link the glideosome to the cytoskeleton to allow parasites to move and invade host cells. Our work establishes that *Plasmodium* GAPMs form a defined, stable heterotrimer, providing a molecular explanation for their long-noted co-dependence and explaining why GAPM proteins often immunoprecipitate with one another^12,14,17^. The same pattern was observed in our mass-spectrometry analyses of both asexual and sexual life-cycle stages in *P. berghei*, and in previous data in several organisms^5,15–17^. In *P. knowlesi*, we see significant enrichment for GAPM3 but not GAPM1. We currently attribute this to the reported poorer solubility of GAPM1^12^, combined with lower sample quantity. We also do not observe higher oligomeric states as previously suggested^12^, which we propose to be an artefact of thermal aggregation (Extended Data Fig. 4C)^21^.

Previous observations in *P. falciparum* have placed GAPM2 as an IMC resident protein that appears during schizont formation^12,22^. Here we use live-cell and ultrastructure imaging in both *P. berghei* and *P. knowlesi*, demonstrating that GAPM2 is a dynamic IMC component. We propose that GAPM2 first appears associated with cytoplasmic vesicles in late rings and trophozoites in *P. berghei* and subsequently redistributes to the parasite periphery, incorporating into the expanding IMC during schizogony. The temporal recruitment of GAPM2 to nascent IMC membranes supports models in which IMC biogenesis is coordinated with MTOC duplication and basal complex progression, ensuring accurate nuclear segregation and daughter cell segmentation^23,24^. This coordination indicates that IMC assembly is tightly regulated with cell cycle progression rather than being a passive consequence of cytoplasmic growth, positioning GAPM2 as a structural scaffold supporting nuclear segregation and IMC expansion.

While IMC development has been extensively studied in *P. falciparum* gametogenesis, much less is known in other *Plasmodium* species^16,23^. In *P. berghei*, earlier ultrastructural studies reported a discontinuous IMC in male gametocytes and an apparent absence in females^25,26^. Consistent with this, we observed rapid GAPM2 recruitment to the surface of activated male gametocytes, where the developing IMC expands discontinuously over the polymerising subpellicular microtubules.

Importantly, our data provide the first evidence of an IMC-like membrane in female gametocytes, appearing immediately after activation as a punctum that matures into an arc and then a ring prior to fertilisation. Ultrastructure imaging using NHS-ester staining revealed no detectable cytoskeletal support at early stages. Although previous studies with IMC marker proteins such as PhIL1, GAP45 and ISPs suggested the absence of IMC in activated females^15,27^, our pull-down analyses detected these proteins within six minutes of activation. This indicates sex-specific differences in IMC composition and cytoskeletal remodelling, consistent with prior observations that microtubule organisation and basal body dynamics differ between male and female gametocytes^28^. These observations suggest that GAPMs represent early structural components of the nascent IMC and may contribute to polarity establishment prior to fertilisation.

Following fertilisation, GAPM2-labelled apical polar rings persisted throughout ookinete differentiation, consistent with early and stable polarity establishment. During elongation, GAPM2 remained enriched at the apical tip and was progressively incorporated into the expanding IMC along the retort, with posterior IMC extension coordinated with basal complex displacement. In mature ookinetes, GAPM2 localised predominantly at the apical end while also distributing uniformly along the IMC, reflecting complete pellicle maturation.

Drawing these results together, we propose that GAPMs form a stable heterotrimeric assembly that is present across the lifecycle of *Plasmodium* parasites. We suggest that this heterotrimer acts as a conserved IMC scaffold whose recruitment is coordinated with nuclear division, cytoskeletal organisation, and polarity establishment throughout parasite development.

### The conservation of the GAPM fold across eukaryotes

We leveraged our structure to explore the conservation of GAPM sequence and fold. Previous sequence-based analyses identified GAPMs as a highly-conserved protein family within Apicomplexa and in non-parasitic Chromerids^12,13,29^. Parasitic apicomplexans appear to always contain a single GAPM3 homologue, but Coccideans (*Toxoplasma*, *Eimeria* and *Neospora*) can have multiple GAPM1 or GAPM2 paralogues. Previous studies in *Toxoplasma* asexual parasites show that GAPM1a and GAPM2a are essential, whereas GAPM1b and GAPM2b are dispensable and much lower expressed^13^. We find that key GAPM1 residues that interact with GAPM2 and GAPM3 in our structure are conserved in *Tg*GAPM1a and *Tg*GAPM1b, with the same true for the GAPM2 homologues. We hence speculate that in organisms with more than one GAPM1 or GAPM2, different stages of the life cycle may contain distinct heterotrimers with each containing one GAPM1 paralogue, one GAPM2 paralogue and GAPM3.

Further to previous analyses, we identify multiple GAPM orthologues in Chromerid species, *Vitrella brassicaformans* and *Chromera velia* (Extended Data Fig. 7A). These orthologues do not classify into the GAPM1/GAPM2/GAPM3 subfamilies and so it is unclear whether they oligomerise in a similar fashion. Chromerids contain other glideosome components, pointing to a semi-conserved architecture^29^. However, it remains to be determined whether these non-invasive organisms use a glideosome for motility in a similar way to apicomplexans.

Interestingly, structural similarity searches using FoldSeek^30^ reveal that the GAPM 6-TM fold shares clear similarity with structural predictions for the FRAG1/DRAM/SFK1 family. These proteins are found in many eukaryotes^31^, with family members primarily associated with maintenance of phospholipid asymmetry and autophagy^32–34^. One member of this family, TMEM150C (also known as DRAM4 and TTN3), has been suggested to form tetramers using equivalent helices to those at the intra-trimeric interface in the GAPM heterotrimer^35^. This raises the possibility of a common oligomerisation mode within similar proteins, although it is unclear whether this reflects true homology or convergent evolution.

### A glideosome model consistent with IMC inter-membrane distance across Apicomplexa

Previous work has defined the glideosome as a set of interacting subcomplexes that crosses the two membranes of the IMC. Structural studies showed that MyoA, MTIP and ELC form the core motor complex that engages actin between the IMC and PM^4,6–9^, while cell biology data demonstrated that GAP40, GAP45 and GAP50 assemble into a complex embedded within the outer membrane of the IMC^11^. GAPM1-3 have been localised to the inner IMC membrane, where they are proposed to link the glideosome to the cytoskeleton^12,13^. Finally, our mass spectrometry data add to the evidence that GAP50 and PhIL1 are proximal to GAPM proteins, with other glideosome components more distal. Integrating our GAPM heterotrimer structure with these data allowed us to propose a unified structural model of the glideosome complex using AlphaFold. Mass spectrometry of our cross-linked samples further supports this model, with proteins expected to be located closer to GAPM2 more highly enriched compared with those predicted to be further away. Notably in *P. berghei* gametocytes, which do not glide, GAPM2 does not coprecipitate with the glideosome motor complex, and the peripheral interactome varies substantially (Fig. 4D). This opens the possibility that GAPMs adopt a role that is distinct from its glideosome association.

The model predicts that the asymmetric surface formed by the GAPM heterotrimer within the IMC lumen engages GAP50. This interface maps to a conserved region on the luminal face of the GAPM complex, with a corresponding conserved surface on GAP50 (Bosch et al., 2012). Notably, this interaction depends on the heterotrimeric arrangement and would not be formed in GAPM homotrimers. This provides a structural basis for previous observations that GAP50 co-immunoprecipitates more strongly with GAPM1 and GAPM2, which contribute to this interface, than with GAPM3 (Bullen et al., 2009).

On the cytoplasmic face, the disordered N- and C-terminal regions of GAPM1-3, which are all unique, contribute to an asymmetric surface, perhaps to interact with proteins that would localise to the cytoplasmic face of the IMC. Mass spectrometry identifies proteins known to localise to this face, including PhIL1, PhIL1-associated proteins, and alveolins, in both *P. berghei* and *P. knowlesi*, with evidence of stage-specific enrichment.

Remarkably, the predicted architecture of the glideosome is consistent with the intermembrane distance within the IMC in several organisms. Electron tomography of *Plasmodium*^36^ and *Toxoplasma*^37^ parasites reveals a highly conserved distance between the inner and outer IMC membranes that closely matches the spacing predicted by our model, approximately 5.5 nm. This intermembrane distance is unusually narrow and appears to be a distinctive feature of the apicomplexan IMC, particularly when compared with other biological membrane systems, such as the endoplasmic reticulum (20-60nm)^38^ or the canonical Gram-negative periplasm (∼25nm)^39^. Whether the glideosome is the key determinant of the intermembrane distance remains to be elucidated.

Taken together, our data show that GAPMs assemble as a stable heterotrimer and are recruited to the IMC in coordination with nuclear division, cytoskeletal organisation, and polarity establishment. We suggest that GAPMs form a part of a stably-maintained scaffold linking the cytoskeleton to the motility apparatus, with peripheral components remodelled in a stage-dependent manner. We speculate that this link will be static rather than transient, enabling the anchoring of the glideosome motor complex to enable motility and invasion.

## Methods

### Ethics statement

All animal procedures were reviewed and approved by the United Kingdom Home Office. Experiments were conducted under UK Home Office Project Licenses 30/3248, PDD2D5182 and PP3589958, in compliance with the *Animals (Scientific Procedures) Act 1986*. Female CD1 outbred mice (6-8 weeks old; Charles River Laboratories) were used in all experiments. The conditions in which mice were kept are a 12 h light and 12 h dark (7 till 7) light cycle, the room temperature is kept between 20 and 24°C and the humidity is kept between 40 and 60%.

### Routine *Pb* culture

To initiate parasite infection, mice were intraperitoneally injected with approximately 1 x 10^6^ blood-stage parasites. Parasite growth was monitored daily by examining thin blood smears stained with Giemsa. The *Plasmodium berghei*-infected blood was collected from female CD1 mice and routinely cultured in RPMI-1640 (Gibco) medium supplemented with 20% FBS (Sigma) and a 1× penicillin/streptomycin (Sigma) antibiotic cocktail. The culture was gassed with a 5% CO₂/O₂/N₂ mixture for 2 minutes and maintained for 8-24 hours at 37 °C on a shaker set at 90 rpm to obtain different erythrocytic stages.

### Generation of transgenic parasite lines: *Pb*GAPM2-GFP

The C-terminus of GAPM2 was endogenously tagged with GFP by single-crossover homologous recombination in *Plasmodium berghei*. To generate the GAPM2-GFP line, a region of the *gapm2* gene downstream of the ATG start codon was amplified using primers T2331 and T2332, ligated into the p277 vector at KpnI/ApaI. The plasmid was linearised using *HindIII* and the cassette was transfected as previously described^15^. A schematic of the endogenous *gapm2* locus (PBANKA_0523900), the targeting construct, and the recombined *gapm2* locus is shown in Extended Data Fig. 1. The oligonucleotides used for mutant line generation are listed in Table SX. For transfections, *P. berghei* ANKA line 2.34 (for GFP tagging) were electroporated as described previously^40^.

To confirm the correct integration of the targeting cassette at the locus was confirmed by the diagnostic PCR using the primer 1 (IntT233) and primer 2 (ol492) (Extended Data Fig. 1C). The list of primers is included in the Supplementary Table 8.

### Purification of schizonts and gametocytes

Five to ten phenylhydrazine pre-treated mice were injected with 1x10^5^-1x10^6^ blood stage parasites. The blood was collected 4 days post infection and cultured for 11 h and 24 h at 37 °C with continuous rotation at 100 rpm. Schizonts were then purified the following day using a 60% (v/v) NycoDenz (Progen) gradient prepared in phosphate-buffered saline (PBS). The NycoDenz stock solution consisted of 27.6% (w/v) NycoDenz in 5 mM Tris-HCl (pH 7.20), 3 mM KCl, and 0.3 mM EDTA. The schizonts were collected from the interface, washed and processed for further analysis.

Gametocyte purification was carried out using a modified version of the protocol described previously^41^. Parasites were first injected into phenylhydrazine-treated mice, and gametocyte production was enriched by sulfadiazine administration 2 days after infection. On day 4 post-infection, blood was collected, and gametocyte-infected erythrocytes were isolated using a 48% (vol/vol) NycoDenz gradient in PBS. The NycoDenz stock solution consisted of 27.6% (wt/vol) NycoDenz in 5 mM Tris-HCl (pH 7.2), 3 mM KCl, and 0.3 mM EDTA. Gametocytes were recovered from the gradient interface, washed thoroughly and activated for further analysis.

### Routine Plasmodium knowlesi culture

*P. knowlesi* parasites were grown *in vitro* in human blood (United Kingdom National Blood Transfusion Service) after confirmation of Duffy-positive (Fy +) blood status (Lorne Blood Grouping Reagents). *P. knowlesi* parasites were maintained at 2% hematocrit in custom-made complete medium, comprised of RPMI-1460 (Invitrogen) with additions including: 2.0 g/L sodium bicarbonate, 2.0 g/L D-glucose, 25 mM HEPES, 0.05 g/L hypoxanthine, 5 g/L AlbuMAX II, 2 mM L-glutamine and 10% (vol/vol) horse serum (Pan Biotech; P30-0702)^42^.

Schizont stages of parasite cultures were enriched using density gradient centrifugation with 60% Percoll (Cytiva) in RPMI-1460, and incubated with 1 uM compound 2 (PKG inhibitor; Blackman lab) for 3 hours to inhibit egress and tighten the synchronisation window for enrichment of late stage schizonts (∼27 hpi specifically)^42^.

### Generation of transgenic parasite lines: *Pk*GAPM2-mNG

Transfectant lines were generated using CRISPR-Cas9 mediated genome editing. Wildtype *Pk*A1-H.1 parasites were co-transfected with: 1) A Cas9-encoding plasmid, containing a guide sequence, targeting the Cas9 enzyme to the C terminus of *Pk*GAPM2, pCas9_GAPM2. 2) Donor DNA, generated by nested PCR, containing the mNG tag flanked by two homology regions to *Pk*GAPM2.

The CRISPR-Cas9 guide plasmid (pCas9_GAPM2, Extended Data Fig. 2B) was designed with a guide sequence targeting Cas9 to the *Pk*GAPM2 (PKA1H_050020900) gene. 20 bp guide sequences, found next to PAM sites (NGG) were identified using EuPaGBT (http://grna.ctegd.uga.edu), and a guide sequence spanning the *Pk*GAPM2 stop codon (CGAAAAATGGTCAATTTTGA) was selected, after first ensuring that there were no off-target binding sites in the *P. knowlesi* genome. Guide sequences were ordered with 15 bp flanking sequences with homology to the pCas9 plasmid vector (ordered sequences in Supplementary Table 8) and ligated into the digested vector as previously described^43^.

Guide plasmids were transformed into competent cells and screened before outgrowth and midi-preps of pCas9_GAPM2.

A nested 3-step PCR approach was used to amplify the homology regions and mNG tag separately, before stitching the constructs together to create the full donor construct, as previously described^43^ (Extended Data Fig. 2). Homology region 1 (HR1) was designed as 774 bp upstream of the stop codon, and HR2 began at the stop codon and was 925 bp. Primers used for donor DNA amplification and expected band sizes from PCRs are outlined in Supplementary Table 8.

For all PCR reactions, CloneAmp HiFi PCR Premix (Takara) was used according to manufacturer’s specifications to a total reaction volume of 25 µL, with 35 cycles of the following conditions: 98 °C for 10 sec, 55 °C for 15 sec, 72 °C for 5 sec/kb of product. For nested PCRs, 1 µL of each PCR product was used as template DNA after PCR clean-up of the original products. The final stitching PCR was ran in a total volume of 50 µL, in 6 independent PCR reactions (total PCR volume of 300 µL) and combined before PCR clean-up (NucleoSpin Gel & PCR Clean-up Mini kit, Macherey-Nagel)^43^. Final PCR products were visualised on a 1% agarose gel to confirm the correct size before precipitating amplicon DNA as previously described^43^.

Late schizont stages (27 hpi) of the *P. knowlesi* wildtype *Pk*A1-H.1 parasite line were harvested and co-transfected with pCas9_GAPM2 and the generated donor DNA using the Amaxa 4D electroporator (Lonza). For transfection, 10 µL of packed late schizonts were mixed with 10 µL mixed DNA containing both the pCas9_GAPM2 plasmid (20 ng) and the donor DNA template (40 ng) with 100 µL P3 Primary Cell electroporation buffer. Schizonts were electroporated using pulse code FP 158 and transferred to 5 mL cultures at 4% hematocrit immediately after. Transfections are kept at routine *P. knowlesi* culture conditions as described above, and 24 hours after transfection, were incubated with 100 nM pyrimethamine for 5 days for positive selection of transfected parasites^43^. Successful transfectant lines were validated using a diagnostic PCR (Extended Data Fig. 2) with primer sequences and expected band sizes in Supplementary Table 8.

### Live-cell imaging: *Pb*GAPM2-GFP

Expression of GAPM2-GFP during blood-stage development was assessed by imaging parasites cultured in schizont medium at different stages of schizogony. Purified gametocytes were examined for GFP signal and localisation at different time points intervals (0-15 min) following activation in ookinete medium. Zygote and ookinete development was tracked over 24 h using the Cy3-conjugated 13.1 antibody, which labels the P28 surface protein. Oocysts and sporozoites were analysed directly in infected mosquito midguts. Imaging was performed using a 63× oil immersion objective on a Zeiss AxioImager M2 microscope fitted with an AxioCam ICc1 digital camera.

### Live-cell imaging: *Pk*GAPM2-MnG

*Pk*GAPM2-MnG was investigated through live cell imaging of schizont stages, purified using density gradient centrifugation and incubated on 1 µM Compound 2 to tighten the level of synchronisation. Schizonts were stained using 1 µg/mL Hoechst-334 to visualise parasite DNA. DNA-stained, purified schizonts were washed and resuspended in complete media to obtain a final concentration of 0.3% purified schizonts. After diluting stained schizonts in complete medium, 100 µl was loaded onto a singular well in the Ibidi µ-Slide VI 0.4. Parasites were imaged using a Nikon Ti E inverted microscope at the 100x oil immersion lens. Images were acquired using NIS-Elements and adjusted in Fiji software (ImageJ 1.54r).

### Western blot analysis for *P. berghei* lysates

Purified schizonts and gametocytes were lysed in lysis buffer containing 10 mM Tris-HCl (pH 7.5), 150 mM NaCl, 0.5 mM EDTA, and 1% NP-40. After addition of Laemmli buffer, samples were heated at 95 °C for 10 min and then centrifuged at 13,000 × g for 5 min. The resulting supernatants were separated on a 4-12% SDS-polyacrylamide gel, and the resolved proteins were transferred onto a nitrocellulose membrane (Amersham Biosciences). Immunoblotting was performed using the Western Breeze Chemiluminescence Anti-Rabbit kit (Invitrogen) with an anti-GFP polyclonal antibody (Invitrogen) diluted 1:1,250, following the manufacturer’s instructions.

### Ultrastructure expansion microscopy (UExM)

Sample preparation of *P. berghei* parasites for UExM was carried out as previously described. Schizont cultures were fixed at 9 and 24 hpi with 2% formaldehyde (FA) to visualise early and mature schizonts, respectively. Purified gametocytes were fixed at 4, 8, and 15 min post-activation in 4% FA to capture distinct developmental stages. Fixed samples were mounted on 12 mm round poly-D-lysine (A3890401, Gibco)-coated coverslips for 30 min and further gels were casted and expanded^44^. Immunolabelling was performed using primary antibodies against α-tubulin (1:1000; Sigma, T9026), GFP (1:250; Thermo Fisher), or centrin (1:500; Thermo Fisher). Secondary antibodies were anti-rabbit Alexa Fluor 488 (Invitorgen) and anti-mouse Alexa Fluor 568 (Invitrogen), each used at 1:1000 dilution. For bulk proteome labelling the gels were stained with Atto 594 NHS or Atto 665 (Merck) diluted to 10 µg/ml diluted in PBS for 90 min at room temperature on a shaker. Images were acquired on a Zeiss Elyra PS.1-LSM780 system and CD7-LSM900, and subsequent image analysis was performed using Zeiss ZEN 2012 (Black edition) and Fiji/ImageJ.

### Structured illumination microscopy

For structured illumination microscopy, 3 µl of schizonts were mixed with Hoechst dye, placed on a microscope slide, and covered with a 50 × 34 mm coverslip to create a thin monolayer and immobilise the cells. Imaging was performed on a Zeiss Elyra PS.1 microscope using structured illumination microscopy (SIM). Cells were scanned with either a Zeiss Plan-Apochromat 63×/1.4 oil immersion objective.

Fluorescence signals were acquired using the following filter sets: BP 420-480 + LP 750 (blue), BP 495-550 + LP 750 (green), and BP 570-620 + LP 750 (red). Z-stacks were collected at 0.2 µm step size. After acquisition, images were processed by maximum intensity projection to merge focal planes. Post-processing and image export were carried out using Zeiss ZEN 2012 (Black edition) software.

### Immunoprecipitation and mass spectrometry

For the interactome analysis of GAPM2-GFP parasites were harvested at the schizont stage (24 hpi), while male gametocytes of the same lines were collected at 6 min post-activation to capture early developmental events. WT-GFP schizonts and gametocytes were included as negative controls for each respective stage.

For *P. knowlesi*, late stage (27 hpi) schizonts were harvested using percoll density gradient centrifugation and compound 2 treatment. Schizonts from the parental *Pk*A1-H.1 parasite line, with no mNG tag were used as negative controls.

Purified parasite pellets were cross-linked with 1% formaldehyde for 10 min to stabilise protein interactions, followed by quenching with 0.125 M glycine for 5 min. Cross-linked samples were then washed three times with PBS (pH 7.5) to remove residual reagents. Protein lysates were prepared and subjected to immunoprecipitation using the GFP-Trap_Agarose kit (Chromotek, gta) following the manufacturer’s protocol. Briefly, the parasite pellets were lysed in lysis buffer (10mM Tris/Cl pH 7.5, 150mM NaCl, 0.5 mM EDTA, 0.5% Nonidet P40) for 30 mins on ice with sonication. The samples are centrifuged at 15,000 xg at 4 °C for 20 minutes and supernatant were added in dilution buffer (10mM Tris/Cl pH 7.5, 150mM NaCl, 0.5 mM EDTA) prior to incubation with GFP-Trap_Agarose beads for 2 h at 4 °C with continuous rotation to capture GFP-tagged proteins and their interacting partners. Unbound proteins were removed by washing in wash buffer (10mM Tris/Cl pH 7.5, 150mM NaCl, 0.05% Nonidet P40), and bead-bound proteins were subsequently digested on-bead with trypsin to generate peptides for mass spectrometry.

### NanoLC-ESI-MS/MS Analysis

Reversed phase chromatography was used to separate tryptic peptides prior to mass spectrometric analysis. Two C18 columns were utilised, an Acclaim PepMap µ-precolumn cartridge 300 µm i.d. x 5 mm 5 μm 100 Å (Thermo Fisher Scientific) and a 75 µm x 25 cm 1.5 µm (Bruker PepSep ULTRA Analytical column). The Ultimate 3000 RSLCnano system (Thermo Fisher Scientific) was used with mobile phase buffer A composed of 0.1% formic acid in water and mobile phase B 0.1 % formic acid in acetonitrile. Samples were loaded on the µ-precolumn equilibrated in 2% aqueous acetonitrile containing 0.1% Trifluoroacetic and peptides were eluted onto the analytical column at 350 nL min^-1^ by increasing the mobile phase B concentration from 4% B to 25% over 36 min, then to 35% B over 10 min, and to 90% B over 3 min, followed by a 10 min re-equilibration at 4% B.

Ultimate 3000 RSLCnano was coupled online to a hybrid timsTOF Pro (Bruker Daltonics, Germany) via a CaptiveSpray nano-electrospray ion source [1]. The timsTOF Pro was operated in Data-Dependent Parallel Accumulation-Serial Fragmentation (PASEF) mode. Peptides were separated by ion mobility depending on their collisional cross sections and charge states. The method settings were as follows: mass range 100 to 1700 m/z, ion mobility range 1/K0 Start 0.6 Vs/cm^2^ End 1.6 Vs/cm^2^, Ramp rate 9.42 Hz and Duty cycle 100%.

### Mass Spectrometry Data Analysis

Raw data was searched using FragPipe version 22.0 (https://fragpipe.nesvilab.org/) against the PlasmoDB database. For the database search, peptides were generated from a tryptic digest with up to two missed cleavages and carbamidomethylation of cysteines as fixed modifications. As variable modifications were added methionine oxidation and acetylation of the protein N-terminus. Scaffold DDA software (https://www.proteomesoftware.com/products/scaffold-dda) and Perseus version 2.0.11.0 (https://maxquant.net/perseus/) were used for data analysis and visualisation of results^45^.

Label-free quantification was performed in MaxQuant using intensity-based absolute quantification (iBAQ). For *P. berghei*, iBAQ values were normalised to the total iBAQ signal per sample to generate relative iBAQ (riBAQ) values. For *P. knowlesi*, iBAQ values were median-normalised across samples prior to downstream comparison.

To quantify GAPM2-associated proteins, ΔriBAQ values were calculated by subtracting the mean riBAQ values of control samples from those of tagged samples (*Pb*GAPM2-GFP − WT-GFP; *Pk*GAPM2-mNG − parental). This subtraction-based approach was used in place of fold-change ratios to provide a linear measure of relative abundance differences between conditions. While ΔriBAQ does not directly report spatial distance, the use of chemical crosslinking prior to lysis supports interpretation of higher ΔriBAQ values as reflecting increased proximity to GAPM2 and/or greater interaction stability within the native complex.

Statistical significance was assessed using two-tailed unpaired Student’s t-tests on biological triplicates. An unpaired design was used because samples were derived from independent parasite cultures with no matched pairing between tagged and control replicates. Multiple hypothesis testing was controlled using the Benjamini-Hochberg procedure with a false discovery rate (FDR) of 5%.

For stage-specific comparisons in P. berghei, ΔriBAQ values were calculated independently for schizont and gametocyte samples (GAPM2-GFP − WT-GFP), and mean ΔriBAQ values were compared between stages using two-tailed unpaired t-tests with Benjamini-Hochberg correction (FDR 5%; see horizontal red dotted line in volcano plots). Volcano plots were generated using the volcanoseR package^46^, with statistical analyses conducted in R (version 4.5.2).

Only proteins identified with a minimum confidence threshold of 95% were considered for downstream analysis. Proteins were only included when more than 1 unique peptides was identified, unless otherwise stated (Supplementary Data 2). All quantitative data and statistical outputs are provided in Supplementary Data 2. Annotations taken from VEuPathDB^47^, with transcriptomics annotations from VEuPathDB^48–50^ and *Plasmo*Gem^51^

### Constructs for recombinant *Pf*GAPM heterotrimer expression

Full-length *Plasmodium falciparum 3D7* GAPM1 (Uniprot ID Q8IE80), GAPM2 (Uniprot ID Q8IFN1) and GAPM3 (Uniprot ID Q8IM26) sequences were codon optimised for expression in *SF9* cells (Integrated DNA Technologies) and synthesised as gene blocks (Integrated DNA Technologies). GAPM1/2/3 were individually cloned into pAcebac1 vector with c-terminal affinity tags. The following constructs were generated: GAPM1 with SGS linker followed by 6 x His tag. GAPM2 with PreScission protease site (Psc) followed by a 2 x strep affinity tag. GAPM3 with DYKDDDDK Flag tag. The resulting constructs were subsequently subcloned into a pBig1 vector, following the protocol described previously^52^.

GAPM core complex lacking the extended N- and C-terminal regions were cloned into individual pAceBac1 vectors with C-terminal affinity tags. Specifically, GAPM1(Δ1-16 and Δ263-303)-6×His, GAPM2(Δ1-43 and Δ265-372)-Twin-Strep, and GAPM3(Δ1-24 and Δ255-284)-DYKDDDDK (FLAG).

### Expression and purification of *Pf*GAPM heterotrimer and *Pf*GAPM core complex in *Sf*9 cells

*Sf9* cells were cultured in serum free - Sf-900^TM^II SFM (1X) (Gibco^TM^) medium at 27 °C with shaking at 120 rpm. Recombinant baculovirus for insect cell expression was made using the Bac-to-Bac system (Invitrogen). Fresh bacmid DNA was transfected into SF9 cells at 0.5 x 10^6^ cells/ml in six-well culture plates using FuGENE HD (Promega) according to the manufacturer’s instructions. After 5 days, 1 ml of the culture supernatant was added to 50ml of 1 x 10^6^ cells/ml and cells were infected for 3 days in a shaking incubator at 27 °C. The P2 virus was isolated by collecting the supernatant after centrifugation at 1500 x g for 15 min and stored at 4 °C. For expression, P2 virus was added at a ratio of 10 mL per 1L of SF9 culture (1.8 to 2.0 x 10^6^ cells/ml) and incubated for 72 hours in a shaking incubator at 27°C. For co-expression of the GAPM1/2/3 core complex, 10 mL of P2 virus corresponding to each individual GAPM construct was combined and used to infect 1 L of Sf9 cells under the same conditions. Cells were harvested by centrifugation at 1500 x g for 15 min at 10 °C. The cell pellet was frozen and stored at −80°C until use.

For purification of the GAPM1/2/3 complex and the GAPM core complex, cell pellet from 4 L *Sf9* expression was resuspended in total of 140 ml of resuspension buffer (20mM HEPES, 150mM NaCl, 10% glycerol, pH 7.4) supplemented with 1 x cOmplete, EDTA-free protease inhibitor cocktail tablet (Roche). The resuspend pellet was sonicated on ice (Amplitude 50, on-time 3 sec, off-time 7 sec, total time – 45 sec). Lysed cells were pelleted by centrifugation at 12,000 x g for 10 min at 10 °C. The supernatant was then transferred into ultracentrifuge tubes and membranes harvested at 146,550 x g for 1 h at 4 °C. Membranes were resuspended in solubilisation buffer (20 mM HEPES, 150 mM NaCl, 10% glycerol, 1% (w/v) DDM (n-Dodecyl β-D- maltoside) (Melford), pH 7.4) and solubilised overnight at 4 °C before centrifugation at 146,550 x g for 30 min at 4 °C to pellet any insoluble material.

For DDM purification, the supernatant containing the solubilised complex was incubated for 2 h at 4 °C with Strep-Tactin®XT 4Flow® high-capacity resin (IBA) equilibrated in wash buffer 1 (20mM HEPES, 150mM NaCl, 10% glycerol, 0.03% DDM, pH 7.4). The resin was washed with 10 column volumes of wash buffer 1. The GAPM complex was eluted with elution buffer (20mM HEPES, 150mM NaCl, 10% glycerol, 0.03% DDM, 50mM biotin, pH 7.4). The eluted sample was concentrated with 30 kDa MW cut-off concentrator (Amicon) and further purified by size-exclusion chromatography on a Superose 6 Increase 10/300 GL column (Cytiva) pre-equilibrated with 20mM HEPES, 150mM NaCl, 0.03% DDM pH 7.4.

For GDN (glyco-diosgenin) purification of the GAPM1/2/3 complex, the supernatant containing the solubilised complex in DDM was incubated for 2 h at 4 °C with Strep-Tactin®XT 4Flow® high-capacity resin (IBA) equilibrated in wash buffer 2 (20mM HEPES, 150mM NaCl, 10% glycerol, 0.03% DDM, 0.03% GDN, pH 7.4) to facilitate detergent exchange. After binding, the resin was washed with 10 CV of wash buffer 2, incubated for 30 min, and subsequently washed with 10 CV of wash buffer 3 (20 mM HEPES, 150 mM NaCl, 10% glycerol, 0.03% GDN, pH 7.4). The GAPM complex was eluted with GDN elution buffer (20mM HEPES, 150mM NaCl, 10% glycerol, 0.03% GDN, 50mM biotin, pH 7.4). The eluted sample was concentrated with 30 kDa MW cut-off concentrator (Amicon) and further purified by size-exclusion chromatography on a Superose 6 Increase 10/300 GL column (Cytiva) pre-equilibrated with 20mM HEPES, 150mM NaCl, 0.02% GDN pH 7.4.

### Western Blot Analysis of recombinant proteins

Proteins were separated by SDS-PAGE and transferred to 0.2 µm PVDF membranes using the Trans-Blot® Turbo™ Transfer System (Bio-Rad, #1704150) with Trans-Blot Turbo™ RTA Mini 0.2 µm PVDF Transfer Kits for 40 blots (Bio-Rad, #1704272) according to the manufacturer’s instructions.

Membranes were blocked in 5% (w/v) skim milk in TBS-T (20 mM Tris-HCl, 150 mM NaCl, 0.1% Tween-20, pH 7.6) for 1 h at room temperature. Blocked membranes were incubated overnight at 4 °C with primary antibodies diluted in blocking buffer. The following primary antibodies were used: mouse anti-6×His tag (Abcam, ab18184), mouse anti-Strep tag (Bio-Rad, MCA2489), and rat anti-FLAG tag (BioLegend, 637301).

After three washes with TBS-T, membranes were incubated for 30 min at room temperature with HRP-conjugated secondary antibodies diluted in blocking buffer. Goat anti-mouse IgG (H+L)-HRP (Abcam, ab47827) and goat anti-rat IgG (H+L)-HRP (Abcam, ab97057) secondary antibodies were used.

Membranes were washed three times with TBS-T and developed using enhanced chemiluminescence (ECL) (Cytiva) reagents according to the manufacturer’s instructions.

### Native mass spectrometry

Purified GAPM complex was concentrated to 5 mg/mL using a 100 kDa molecular-weight-cutoff centrifugal concentrator (Amicon Ultra) and buffer-exchanged into 200 mM ammonium acetate (pH 8.0) supplemented with 0.02% DDM using a Bio-Spin 6 desalting column (Bio-Rad). The protein was then diluted into the final native MS buffer consisting of 200 mM ammonium acetate and G1-OGD at 2× its critical micelle concentration and introduced into the mass spectrometer using in-house-prepared gold-coated borosilicate capillaries. Spectra were acquired on a Q Exactive UHMR mass spectrometer (Thermo Fisher Scientific) using the following settings: 1.2 kV capillary voltage, 100% S-lens RF, quadrupole selection of 1,000-20,000 m/z, 200 V collisional activation in the HCD cell, trapping gas pressure setting 7.5, 200 °C source temperature, and 12,500 resolving power. The noise threshold was set to 3 (rather than the default 4.64), and no in-source dissociation was applied. Data were processed in Xcalibur 4.2 (Thermo Fisher Scientific).

### Cryo-EM sample preparation and data collection

For cryo-EM grid preparation, 3 µL of 3mg/ml of purified GAPM, prepared in either DDM or GDN, protein complex was applied onto glow-discharged holey carbon grids (Quantifoil R1.2/1.3 Cu, 300 mesh). The grids were blotted using a Vitrobot Mark IV (ThermoFisher) at 100% humidity and 4°C, blotted for 3 sec at blot force 3 before being plunged into liquid ethane.

The first DDM dataset (hereafter referred as preliminary dataset) was collected using a ThermoFisher Titan Krios (300 kV) equipped with a K3 detector and energy filter (20-eV slit size) (Gatan) using automated collection (ThermoFisher EPU). Movies were acquired at 105,000 x magnification (0.83 Å/pixel, 2.4 sec exposure, 40 frames, total electron dose of 42 e^-^/A^2^, -1 to -2.4 μm defocus range) resulting in total of 13,316 movies.

The second DDM dataset (hereafter referred as dataset 2) was collected using a ThermoFisher Titan Krios (300 kV) equipped with a K3 detector and energy filter (20-eV slit size) (Gatan) using multi-grid automated collection with AFIS (ThermoFisher EPU) over two grids. Movies were acquired at 105,000 x magnification (0.83 Å/pixel, 2.4 sec exposure, 40 frames, total electron dose of 39.5 e^-^/ Å ^2^, -1 to -2.4 μm defocus range) resulting in total of 35,306 movies.

The GDN dataset was collected using a ThermoFisher Titan Krios (300 kV) equipped with a K3 detector and energy filter (20-eV slit size) (Gatan) using automated collection (ThermoFisher EPU).Movies were acquired at 105,000 x magnification (0.83 Å/pixel, 2.4 sec exposure, 40 frames, total electron dose of 40.5 e^-^/A^2^, -1 to -2.2 μm defocus range) resulting in total of 13,836 movies.

### Cryo-EM data processing of GAPMs in DDM

Data processing was done in cryoSPARC45 v4.7.1^53^, unless otherwise specified, with the overall workflow summarised in Extended Data Fig 4. Standard 2D classification was performed using customised parameters (total number of online EM iterations = 25; batchsize per class = 400), unless stated otherwise.

For both, preliminary dataset and dataset 2, raw movies were processed using patch motion correction (multi). The Contrast Transfer Function (CTF) was estimated by patch CTF estimation (multi). Data was split into 68 exposure groups before selecting high-quality images were selected for further processing based on defocus ranges, astigmatism, CTF-fit resolution, ice thickness and total full-frame motion distance.

Initial 2D templates were generated from a subset of 300 micrographs in the preliminary dataset. Particles were blob-picked and extracted with 4× binning (corresponding to a pixel size of 3.32 Å) (82,748 particles), followed by two rounds of 2D classification. Well-defined classes were used to train a Topaz model on the same subset of micrographs, which was subsequently applied to the full dataset for Topaz particle picking (2,083,286 particles). Two additional rounds of 2D classification were performed, and the best-resolved particle classes (565,540 particles) were imported into RELION 5.0^54^ for an additional round of 2D classification. Low-quality classes were discarded, and the refined particle set of 276,748 particles was re-imported into cryoSPARC for downstream analysis.

The resulting 2D classes were used as templates for particle picking in dataset 2. A total of 10,704,382 particles (4× binned, pixel size = 3.32 Å) were extracted and subjected to two rounds of 2D classification, yielding 3,482,153 selected particles. From these, 1,087,944 particles from the best-resolved classes were used for ab initio reconstruction (C1 symmetry, three classes). The resulting volumes were used as reference maps for heterogeneous refinement of the full 3,482,153-particle dataset. One class exhibited superior density compared to the others, while another displayed apparent dimeric rather than trimeric organisation.

A total of 1,252,695 particles from the best class were re-extracted without binning and further processed using the HR-HAIR pipeline^55^ with minor parameter adjustments. One round of 2D classification (*Number of 2D classes - 200; Maximum resolution - 3 Å; Initial classification uncertainty factor – 1; Number of final full iterations - 20; Number of online-EM iterations - 80; Batchsize per class - 400*) was performed, and high-quality classes exhibiting clear secondary structure features were selected (165,262) for a subsequent ab initio reconstruction *(Number of Ab-Initio classes – 2; Maximum resolution - 3 Å; Initial resolution - 5 Å; Fourier radius step - 0.005; Initial minibatch size – 300; Final minibatch size – 1000)*.

The best-resolved class (98,384 particles) was locally refined (*Standard deviation of prior over rotation – 2°; Standard deviation of prior over shifts – 1 Å; Rotation search extent - 2°; Shift search extent – 1 Å; Initial lowpass resolution – 6 Å, Re-center rotations each iteration? – yes; Re-center shifts each iteration? - yes*) using reference map obtained from reconstruct only and a soft mask excluding micelle density, following local CTF correction. The resulting map was sharpened using the RELION 5.0 post-processing. The final reconstruction reached a global resolution of 3.4 Å, based on the FSC = 0.143 criterion.

### Cryo-EM data processing of GAPMs in GDN

Data processing was done in cryoSPARC45 v4.7.1(Punjani et al., 2017), unless otherwise specified, with the overall workflow summarised in Extended Data Fig 5. Standard 2D classification was performed using customised parameters (total number of online EM iterations = 25; batchsize per class = 400), unless stated otherwise.

Raw movies were processed using patch motion correction (multi). The Contrast Transfer Function (CTF) was estimated by patch CTF estimation (multi). Data was split into 70 exposure groups before selecting high-quality images were selected for further processing based on defocus ranges, astigmatism, CTF-fit resolution, ice thickness and total full-frame motion distance.

Particles were blob-picked and extracted with 4× binning (corresponding to a pixel size of 3.32 Å) (5,502,206 particles), followed by two rounds of 2D classification yielding 2,253,220 selected particles. The particle set was subjected to ab-initio reconstruction (three maps, C1 symmetry). The resulting volumes were used as reference maps for heterogeneous refinement. One class exhibited superior density compared to the others and was selected for further processing (973,566 particles).

These 973,566 particles were then re-extracted at full resolution (without binning) and further processed using the HR-HAIR pipeline(Kim et al., 2025), with minor parameter adjustments.

One round of 2D classification (*Number of 2D classes - 200; Maximum resolution - 3 Å; Initial classification uncertainty factor – 1; Number of final full iterations - 20; Number of online-EM iterations - 80; Batchsize per class - 400*) was performed, and high-quality classes exhibiting clear secondary structure features were selected for a subsequent ab initio reconstruction (504,885 particles) *(Number of Ab-Initio classes – 3; Maximum resolution - 3 Å; Initial resolution - 5 Å; Fourier radius step - 0.005; Initial minibatch size – 300; Final minibatch size – 1000).* The best-resolved class was locally refined (*Standard deviation of prior over rotation – 2°; Standard deviation of prior over shifts – 1 Å; Rotation search extent - 2°; Shift search extent – 1 Å; Initial lowpass resolution – 6 Å, Re-center rotations each iteration? – yes; Re-center shifts each iteration? - yes*) using reference map obtained from reconstruct only and a soft mask excluding micelle density. The resulting map was sharpened using the Relion 5.0 post-processing. The final reconstruction reached a global resolution of 3.6 Å, based on the FSC = 0.143 criterion.

### Model building

The amino acid sequences of *P. falciparum* GAPM1, GAPM2, and GAPM3 were imported into ModelAngelo^56^ (implemented in RELION 5.0) for initial model building. The resulting model was manually inspected, and regions corresponding to missing or poorly defined density were manually built and subsequently refined through iterative manual adjustments in COOT ^57^ and real space refinement in Phenix^58^.

Structural analyses were carried out in ChimeraX ^59^, using the interface command to identify interacting residues. The matchmaker command was used to compare GAPM1, GAPM2 and GAPM3, with RMSD for the pruned atom set, corresponding to the TM regions (RMSD_TM_) reported. RMSD_TM_ for GAPM1:GAPM2 = 0.93 Å; GAPM2:GAPM3 = 0.95 Å; GAPM1:GAPM3 = 1.0 Å. RMSD of all atoms built for GAPM1:GAPM2 = 3.5 Å; GAPM2:GAPM3 = 2.0 Å; GAPM1:GAPM3 = 5.6 Å.

### Bioinformatics

Structure-based homology searches were carried out using FoldSeek^30^, using GAPM2 from the CryoEM structure as input. Sequence-based homology searches were carried out using JACKHMMER^60^. For conservation analysis, representative species were curated for each GAPM paralogue to avoid over-representation, and aligned using Clustal Omega^61^. Each alignment was then provided as input to ConSurf^62^ to calculate per-residue conservation scores. To visualise surface conservation, scores were rendered onto the CryoEM heterotrimer structure. The standard ConSurf colourblind-friendly colour scheme was used for visualisation.

### AlphaFold

Structure predictions were performed using a local installation of AlphaFold2-Multimer^63^, or AlphaFold3^64^ for predictions that were too large for AlphaFold2, all using default settings. In each case, the model with the highest iPTM score was used. The glideosome AlphaFold model was predicted as three subcomplexes: i) GAPM1:GAPM2:GAPM3:GAP50; ii) GAP50:GAP40:GAP45; iii) GAP40:GAP45:ELC:MTIP:MyoA. To assemble an overall model, the three subcomplexes were combined in ChimeraX, using the matchmaker command to align the overlapping subunits. The GAPM heterotrimer cryo-EM structure was also superposed with the subcomplex i). This glideosome model was adjusted in ChimeraX so that the two transmembrane regions were positioned parallel to one another by rotating around the hinge between the GAP50 globular domain and its C terminal helix. PAE plots were all generated in ChimeraX.

## Data accessibility

GAPM heterotrimer structure data available at wwPDB (PDB 28NA, EMDB EMD-56647 and EMD-56677). GAPM2 interactome Mass Spectrometry Data available at PRIDE (XXXX and YYYYY). Other data available on request from authors.

## Acknowledgements

We thank A.A Holder, B. Cooper, and members of the Lau Lab for comments on the manuscript. We thank R. Matadeen, F. Moreira-Leita, C. Mycroft-West and E. Lowe at the COSMIC cryo-EM facility (University of Oxford) for support with data collection and data processing, as well as M. Pitt and B. Rozman for COSMIC cluster maintenance. We thank Cleidiane Zampronio at Warwick University for mass spectrometry methods and Bio Support Unit, University of Nottingham for maintenance of mice used in this study.

## Author contributions

A.M., G.R., R.T. and C.K.L. conceived and designed the study. A.M. and R.T. generated the *Pb*GAPM2-GFP parasites. A.M. R.T. D.B. and M.Z. performed the *P. berghei* live cell and U-ExM microscopy, the mouse-related work, and sample preparation for mass spectrometry. A.I. and R.W.M generated the *Pk*GAPM2-mNG parasites, performed the *P. knowlesi* live cell microscopy, and sample preparation for mass spectrometry. A.B. collected mass spectrometry data, and E.C.T. analysed the mass spectrometry data. G.R. expressed, purified and characterised the *Pf*GAPM complex and collected the cryo-EM data. G.R. processed the cryo-EM data with some support from C.K.L., then G.R. and C.K.L. built and refined the structure. C.K.L. performed the bioinformatics and structural modelling with assistance from G.R.. D.Z. and J.R.B. collected and analysed the recombinant native mass spectrometry data. A.M., G.R., R.T. and C.K.L. contributed to the initial draft, with C.K.L. and G.R. preparing the manuscript with comments and figures from R.T., A.M., E.C.T., A.I. and R.W.M.

---

**Extended Data Figure 1.**
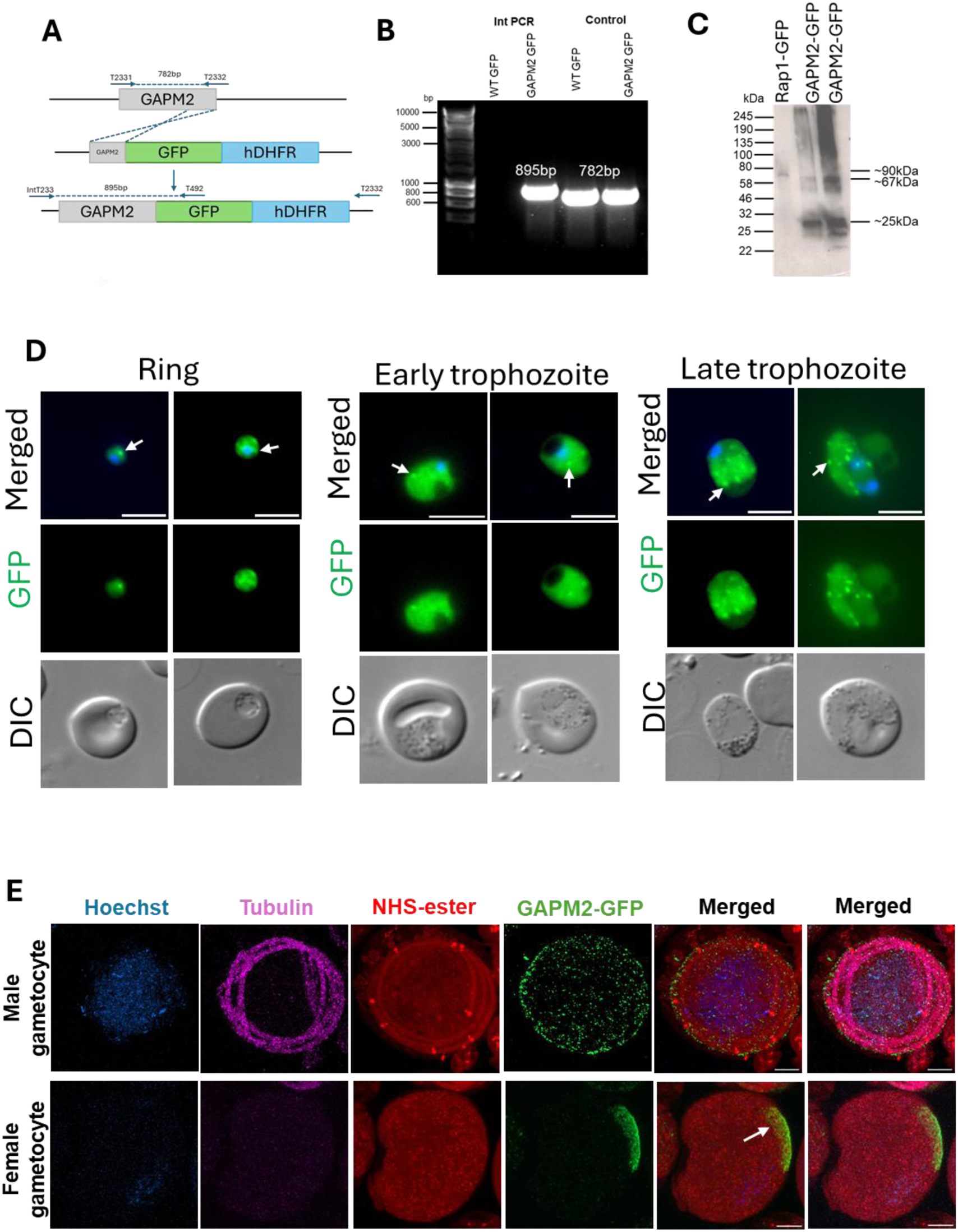
Generation, validation, and subcellular localization of *Pb*GAPM2-GFP in *Plasmodium berghei.* **(A)** Construct design diagram detailing the insertion of *Pb*GAPM2-GFP by homologous recombination. **(B)** PCR verification of successful insertion of *Pb*GAPM2-GFP, as well as a GFP-only control. **(C)** Western Blot analysis showing expression of GFP and *Pb*GAPM2-GFP. **(D)** Further representative images of asexual ring and trophozoite stages in *Pb*GAPM2-GFP, showing punctate vesicular distribution pattern (arrows). **(E)** Ultrastructure expansion microscopy (U-ExM) of 8 min post-activation gametocytes co-stained with anti-GFP (green, arrow), anti-tubulin (magenta) to visualize microtubules, and NHS ester (red) to broadly label cellular ultrastructure to assess the spatial relationship between GAPM2-enriched IMC membranes and basal bodies. Scale bars 5 µm.

**Extended Data Figure 2.**
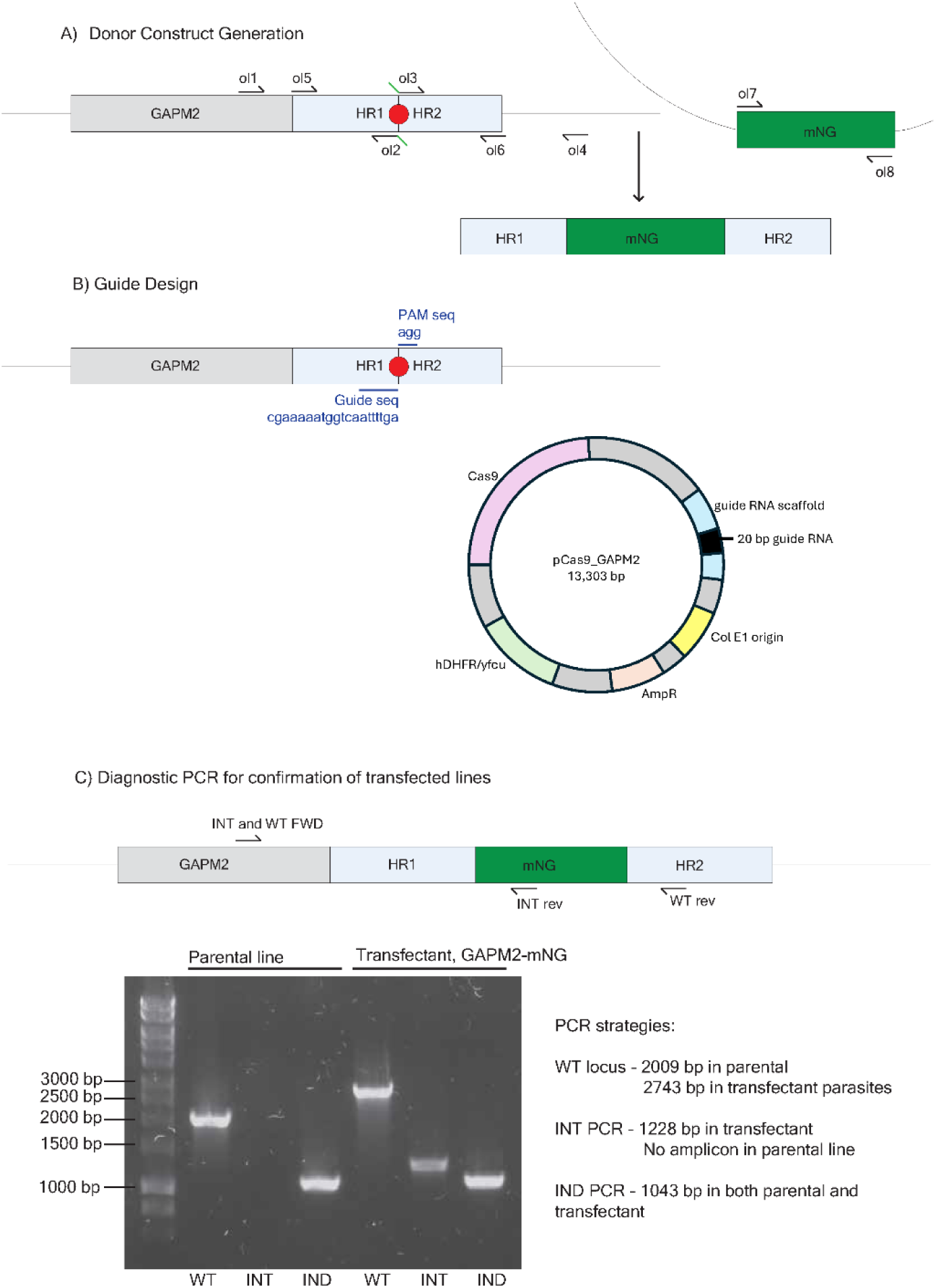
Generation of DNA constructs to introduce an mNG tag at the C terminus of *Pk*GAPM2. **(A)** The final donor DNA construct, with mNG flanked between two homology regions (HR1-mNG-HR2), was generated by a three-step PCR approach, with homology regions (HR1 and HR2) and mNG amplified in 3 PCR reactions. PCR products were combined and sequentially stitched together to generate the final construct. **(B)** A guide sequence, adjacent to a PAM motif, and spanning the *pkgapm2* stop codon, was selected and cloned into a pCas9 vector containing expression cassettes for the Cas9 endonuclease (Cas9), a selection marker cassette encoding pyrimethamine resistance (hDHFR) and an ampicillin resistance cassette (AmpR). **(C)** Diagnostic PCRs confirming integration of mNG into the *Pk*GAPM2-mNG line. PCRs amplifying the wild-type parental locus (WT), the integrant locus (INT), and an independent locus (IND) were performed on both the parental and transfected line, demonstrating a positive integration in only the transfectant *Pk*GAPM2-mNG line. Expected band sizes for each PCR strategy are listed. All primer sequences used in PCRs are listed in Supplementary Table 8.

**Extended Data Figure 3.**
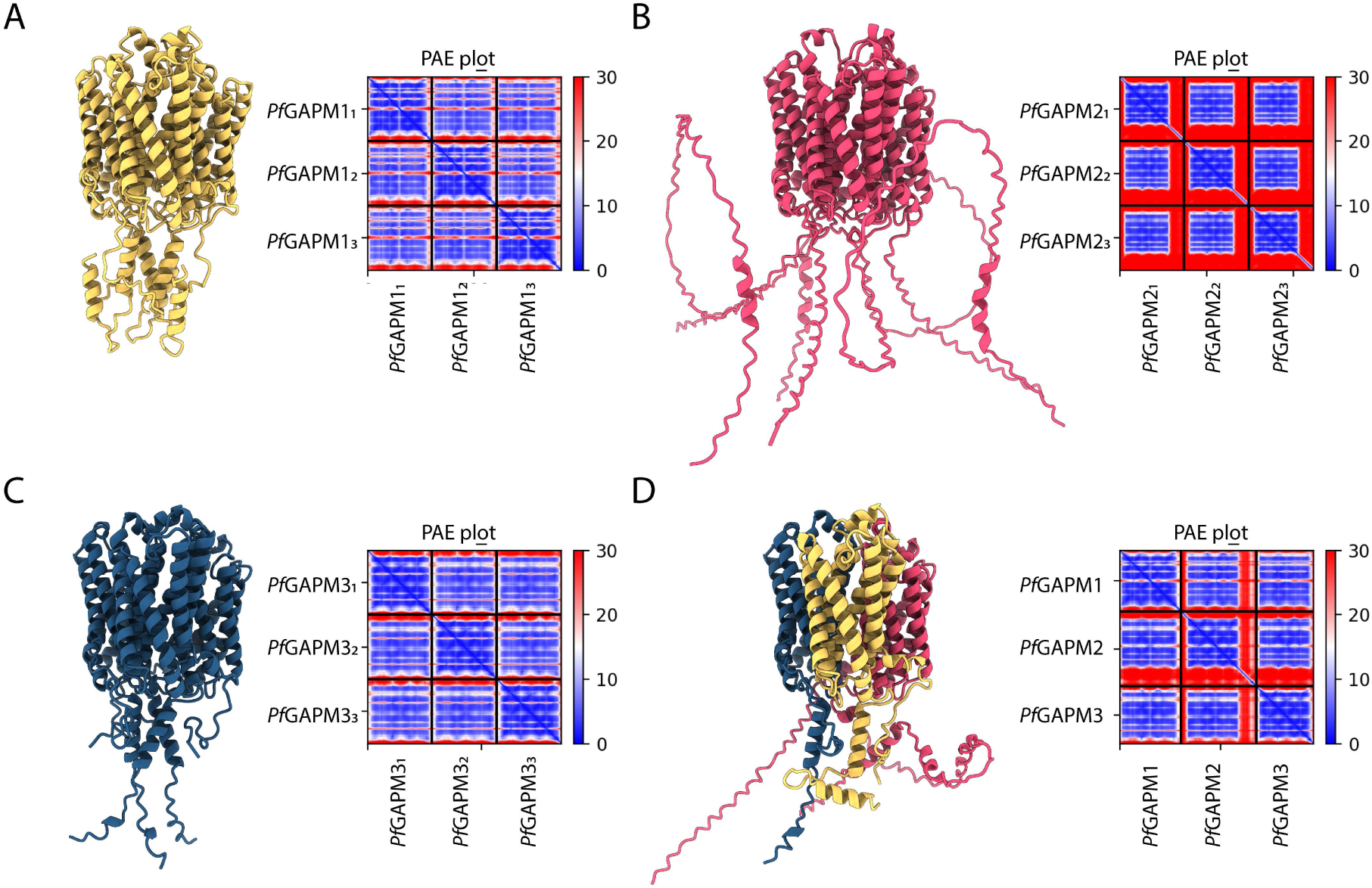
AlphaFold2 predicts GAPM complexes assemble as both homo-and heterotrimers with comparable confidence. **(A)** AlphaFold2 model of the *Pf*GAPM1 homotrimer (UniProt ID: Q8IE80) with the corresponding Predicted Aligned Error (PAE) plot. **(B)** AlphaFold2 model of the *Pf*GAPM2 homotrimer (UniProt ID: Q8IFN1) with the corresponding PAE plot. **(C)** AlphaFold2 model of the *Pf*GAPM3 homotrimer (UniProt ID: Q8IM26) with the corresponding PAE plot. **(D)** AlphaFold2 model of the *Pf*GAPM1/2/3 heterotrimer with the corresponding PAE plot. For all models, the PAE colour scale (blue to red) indicates the expected positional error (Å), with lower values (blue) reflecting higher confidence in the relative positioning of residues.

**Extended Data Figure 4.**
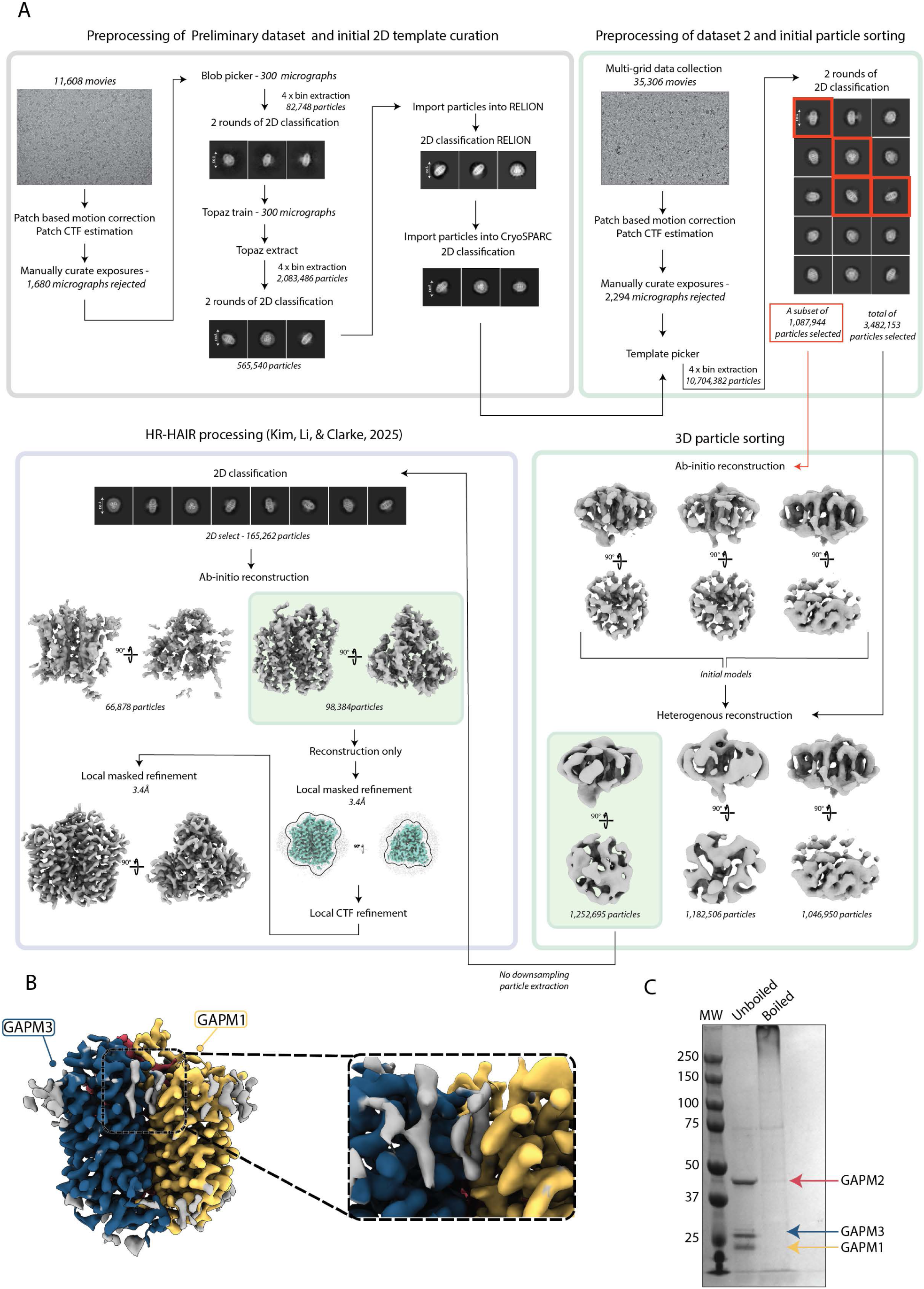
Cryo-EM processing workflow for *Pf*GAPM heterotrimer in DDM and lipid-associated density in the GAPM heterotrimer. **(A)** Cryo-EM data processing workflow for the *Pf*GAPM heterotrimeric complex purified in DDM detergent. All image processing was performed in CryoSPARC v4, with HR-HAIR processing carried out as described by Kim, Li, and Clarke (2025). **(B)** Cryo-EM density map of the *Pf*GAPM heterotrimer coloured by subunit (GAPM3, blue; GAPM2, red; GAPM1, yellow), revealing additional lipid-associated density (grey) located at the interface between the GAPM1 and GAPM3 subunits. **(C)** SDS-PAGE analysis of *Pf*GAPM samples under boiled (95 °C, 10 min) and unboiled conditions, showing that boiling causes the complex to migrate as a high-molecular weight species (>250 kDa).

**Extended Data Figure 5.**
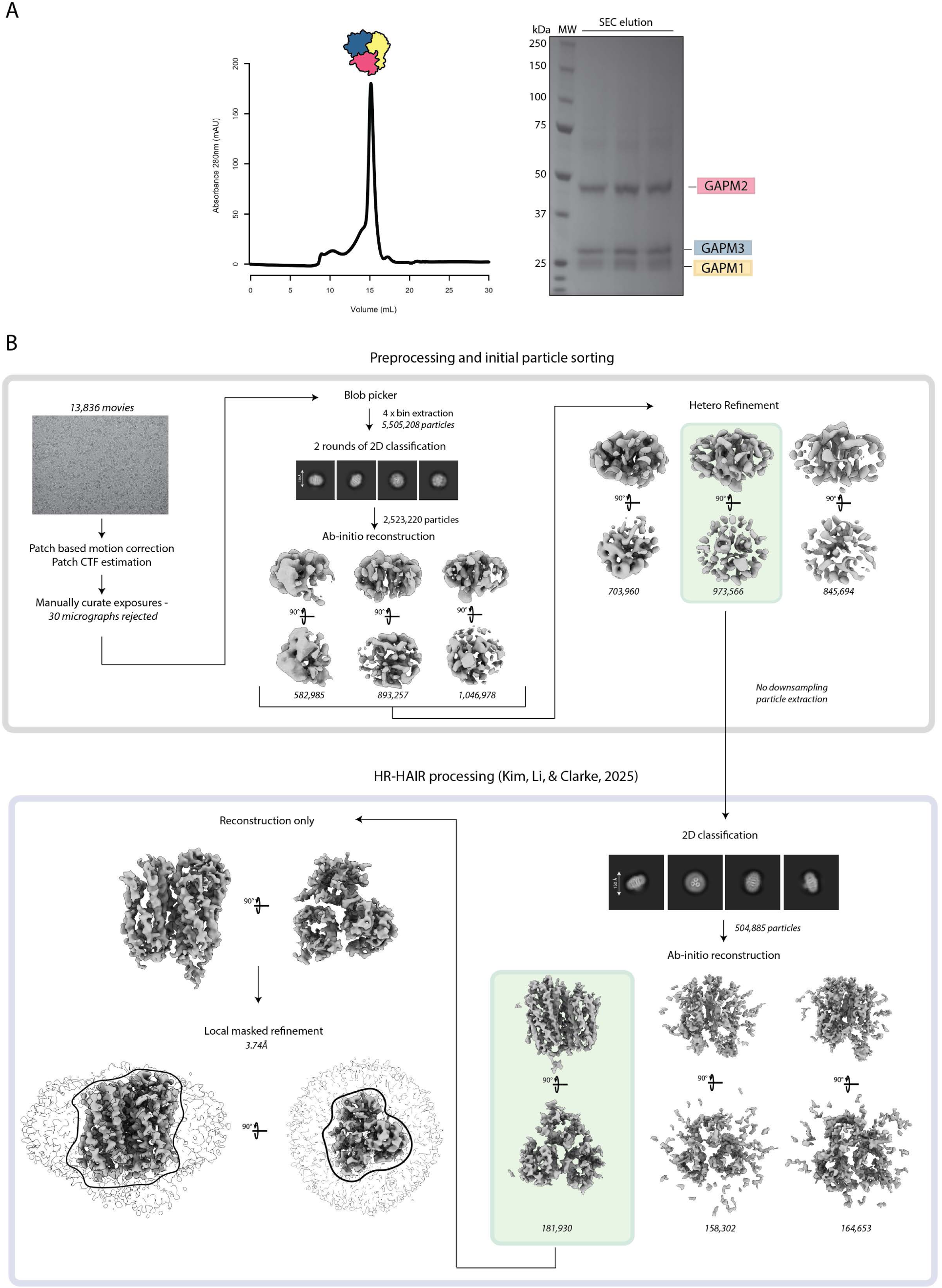
Purification and Cryo-EM processing for *Pf*GAPM heterotrimer in GDN. **(A)** SEC profile and SDS PAGE gel showing purification of *Pf*GAPM1/2/3 in GDN detergent. **(B)** Cryo-EM data processing workflow for the *Pf*GAPM heterotrimeric complex purified in GDN detergent. All image processing was performed in CryoSPARC v4, with HR-HAIR processing carried out as described by Kim, Li, and Clarke (2025).

**Extended Data Figure 6.**
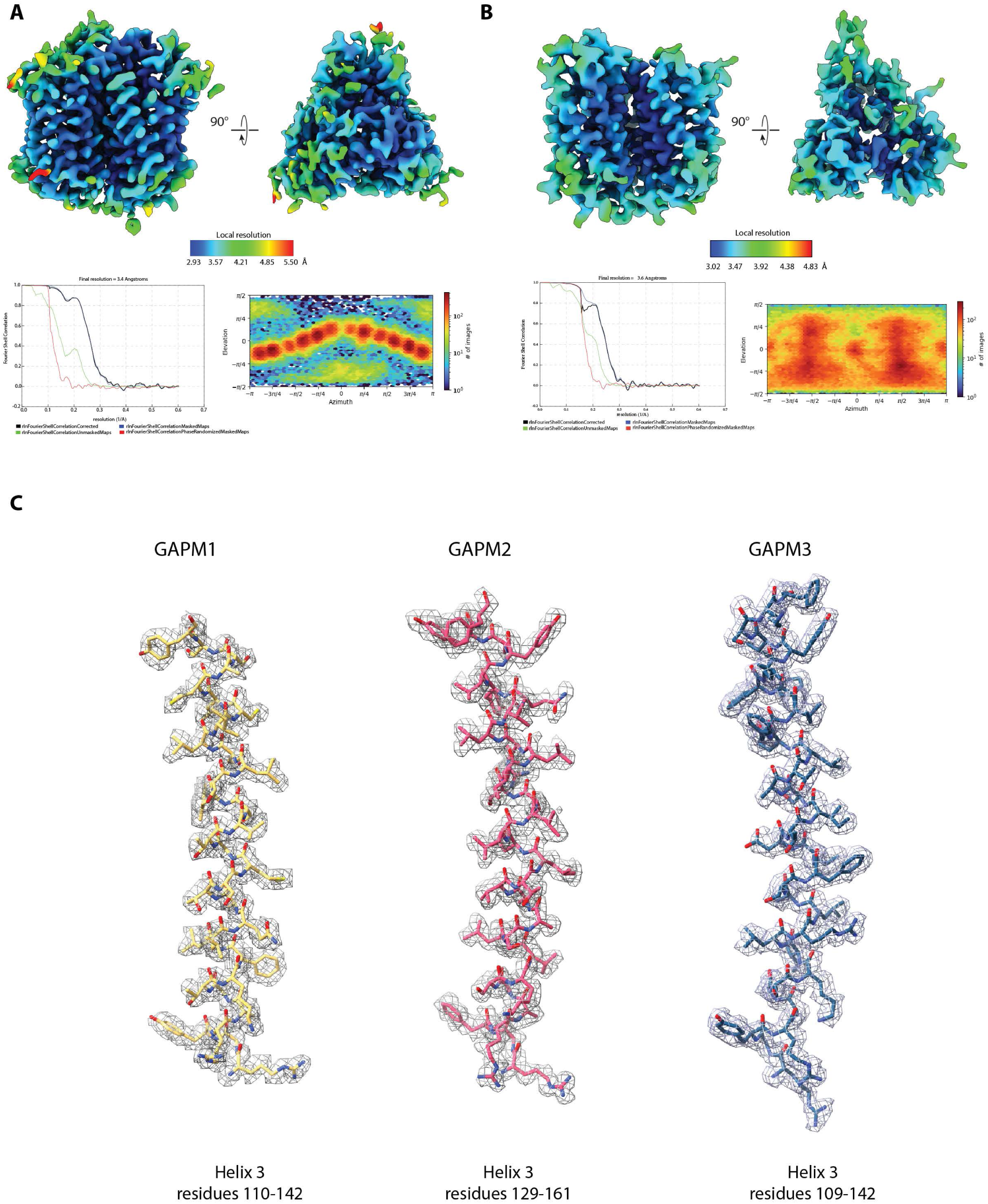
Cryo-EM SPA validation of *Pf*GAPM1/2/3 density maps. **A, B)** Post-processed cryo-EM density maps of the *Pf*GAPM heterotrimer in DDM (A) and GDN (B) detergents, coloured by local resolution (Å) with the corresponding colour keys shown below each map. Fourier shell correlation (FSC) curves are shown for both datasets, with global resolutions determined using the gold-standard 0.143 criterion. The angular distribution plot from CryoSPARC is displayed, illustrating the number of particles contributing to each viewing direction). **(C)** Refined *Pf*GAPM1/2/3 model highlighting helix 3 in all three subunits. The cryo-EM density (grey mesh) is contoured at 1σ, and the model is shown in ball-and-stick representation, coloured yellow (GAPM1), dark pink (GAPM2), and dark blue (GAPM3).

**Extended Data Figure 7.**
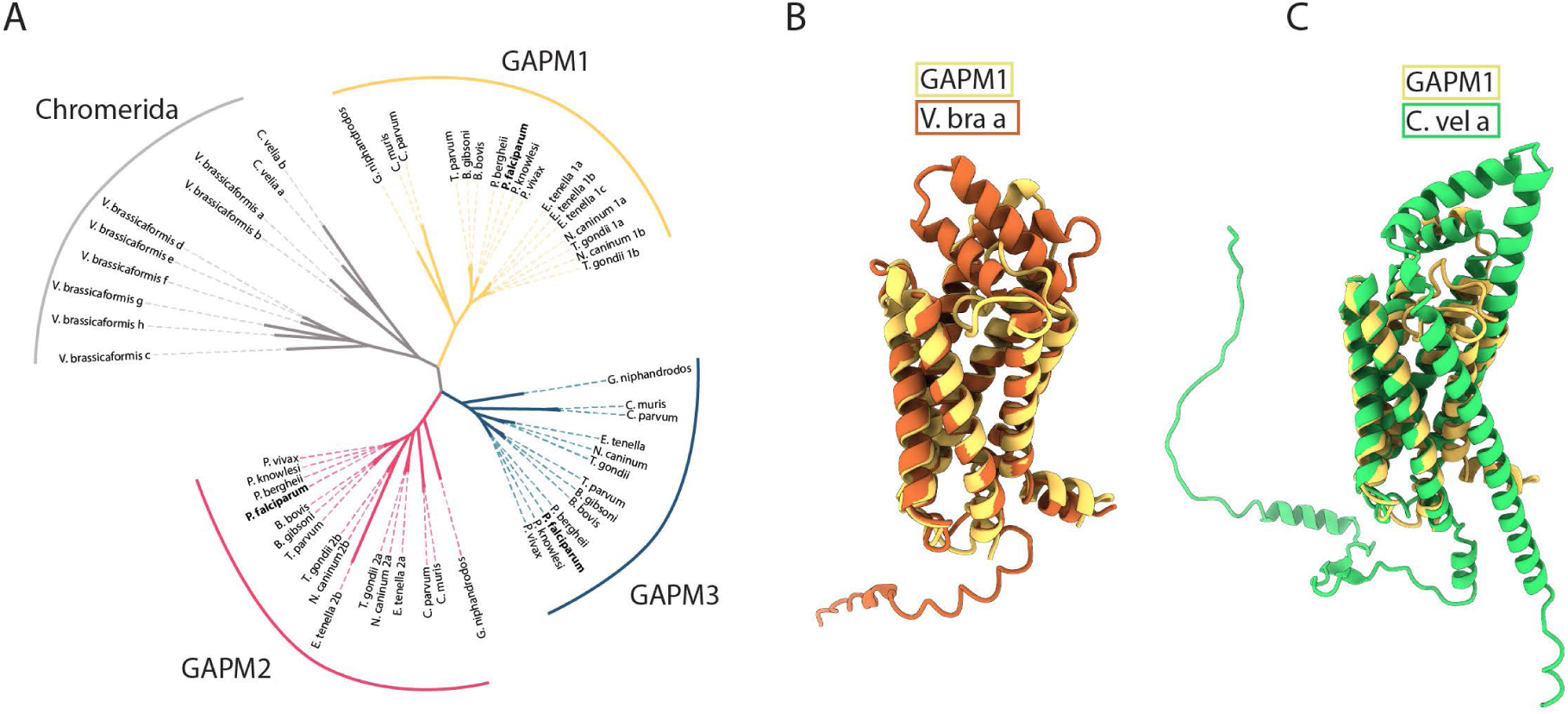
Sequence conservation and structural homology of the GAPM protein family. **(A)** Phylogenetic tree based on sequence homology of representative GAPM proteins across Apicomplexa and Chromerida. **(B)** Structural superposition of the experimentally determined *Pf*GAPM1 structure with the AlphaFold prediction of *Vitrella brassicaformis* GAPM1. **(C)** Structural superposition of the experimentally determined *Pf*GAPM1 structure with the AlphaFold prediction of *Chromera velia* GAPM1.

**Extended Data Figure 8.**
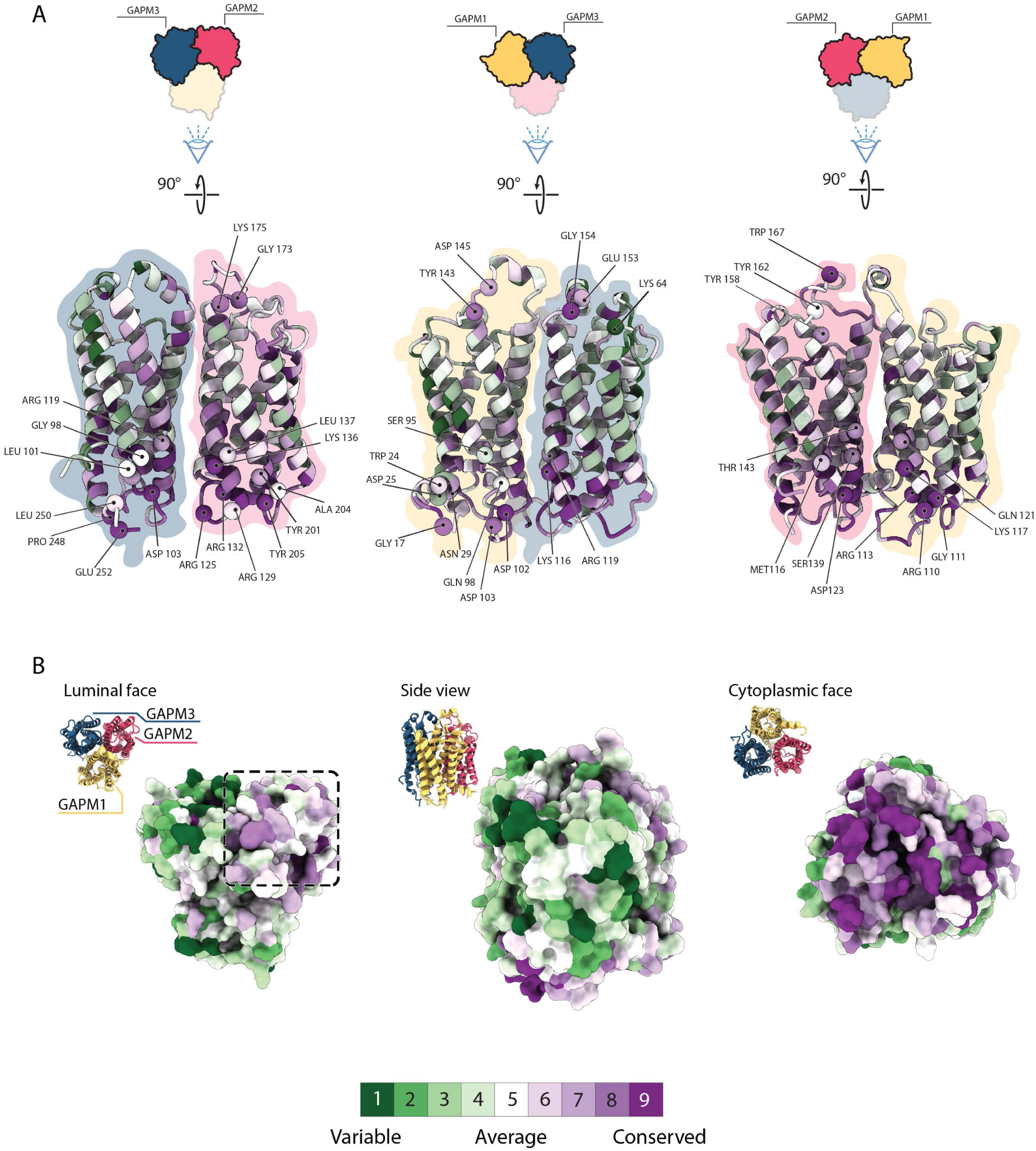
Conservation analysis of *Pf*GAPM1/2/3 complex. **(A)** Residue conservation at the interaction interfaces of the *Pf*GAPM1/2/3 heterotrimer. Shown are the interfaces formed by GAPM1-GAPM2 with GAPM3, GAPM1-GAPM3 with GAPM2, and GAPM2-GAPM3 with GAPM1. Spheres mark Cα positions of residues engaged in hydrophilic interactions (hydrogen bonds and salt bridges) with the neighbouring subunit and are coloured according to their ConSurf-derived conservation scores (colour scale shown below). Cartoon top-view representations of the GAPM1/2/3 complex are provided for orientation. **(B)** Overall conservation of the GAPM1/2/3 heterotrimer. The luminal face of the GAPM2 subunit shows a conserved patch (dashed box), whereas the cytoplasmic face high conservation. Conservation scores were obtained from ConSurf analysis (colour scale shown below), using alignments for each *Pf*GAPM comparing between diverse apicomplexan species (Supplementary Data 5,6,7).

**Extended Data Figure 9.**
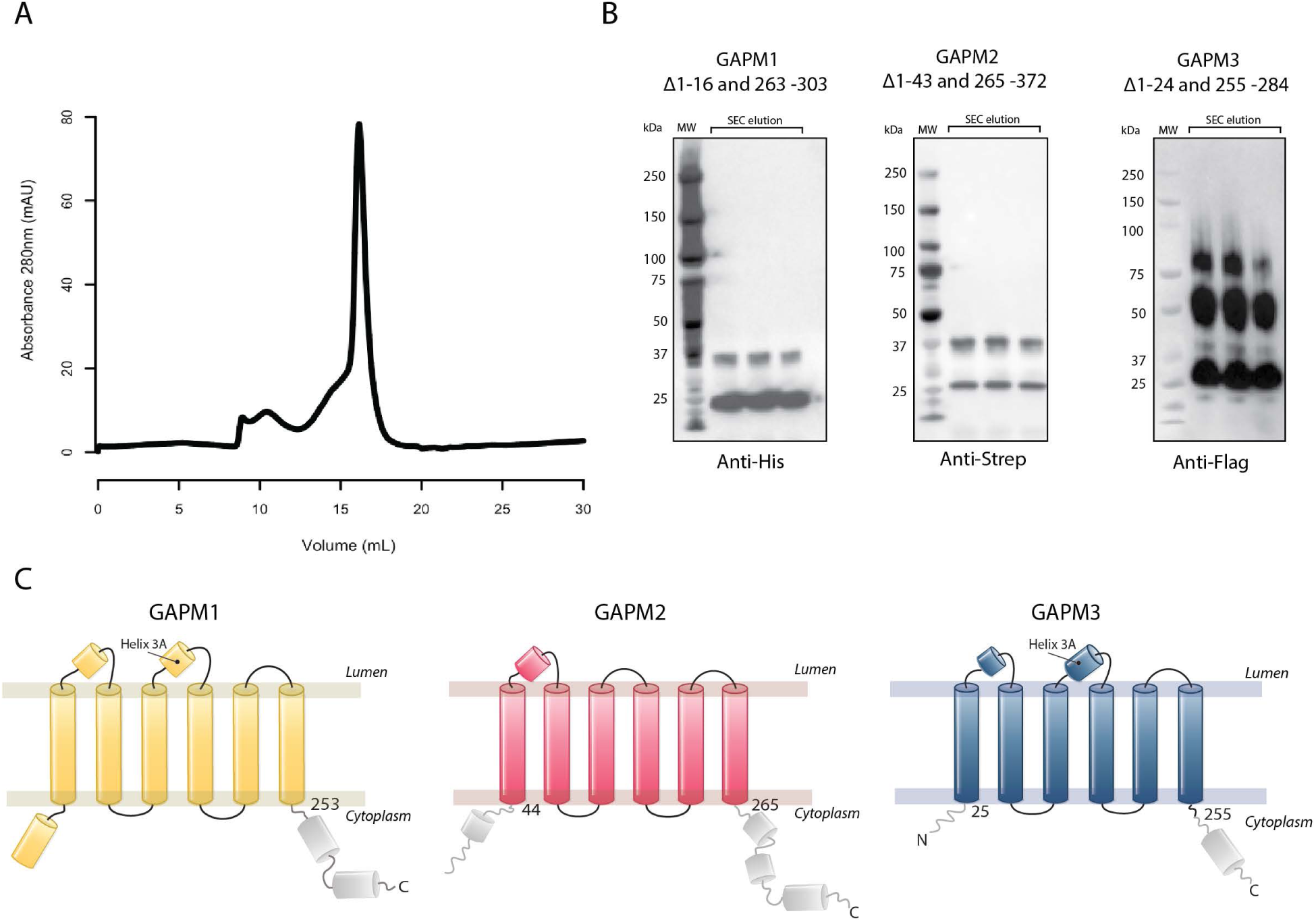
Purification of the *Pf*GAPM heterotrimer core complex. **(A)** SEC profile and **(B)** Western blots for the GAPM heterotrimer core complex showing that all GAPMs are preset in the SEC peak at ∼16 ml, consistent with a heterotrimeric complex. **(C)** Topology diagrams of *Pf*GAPM1 (residues 17-253), *Pf*GAPM2 (residues 44-265), and *Pf*GAPM3 (residues 25-255). Grey segments indicate regions not resolved in the cryo-EM map and removed from the resulting constructs.

**Extended Data Figure 10.**
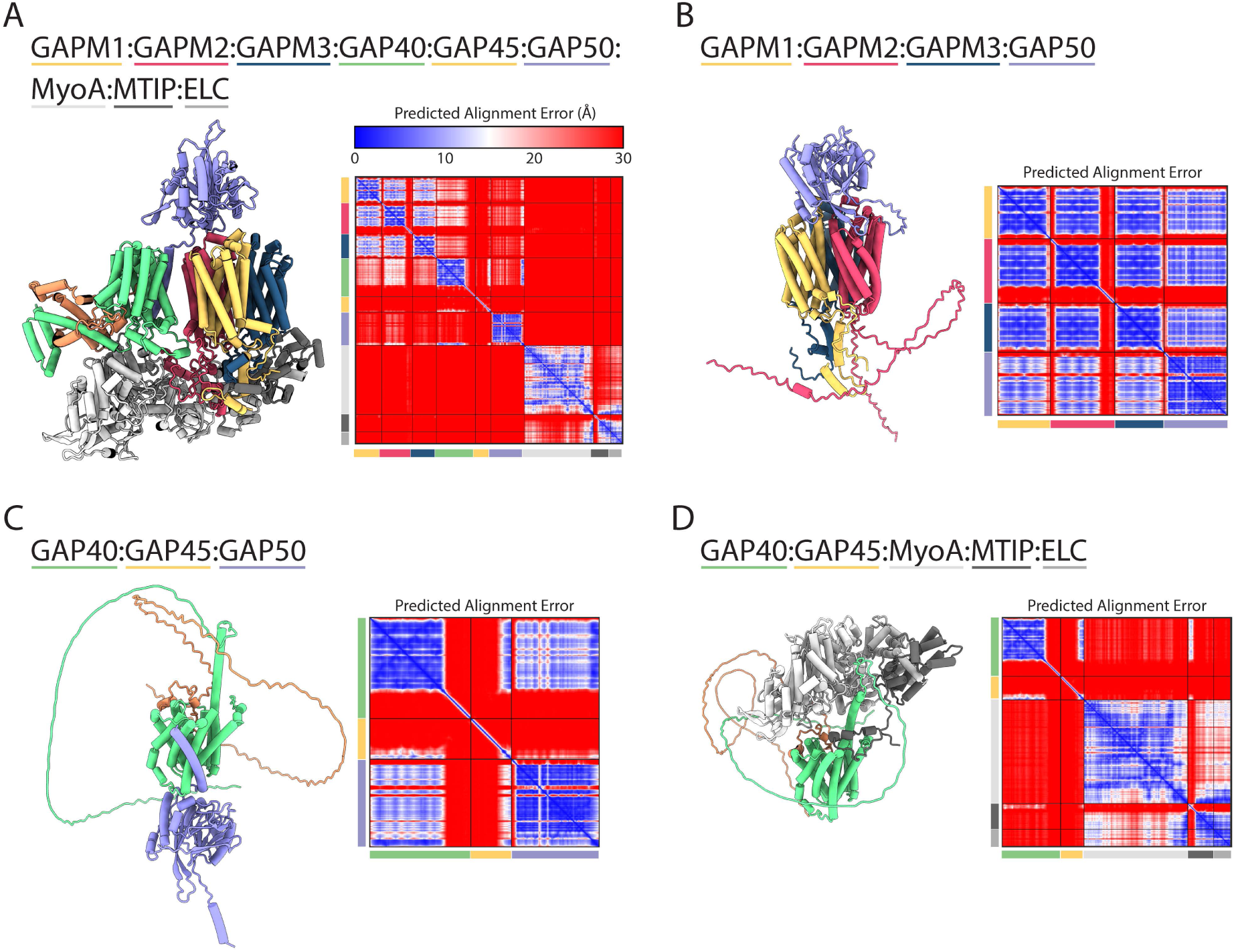
AlphaFold2 predictions of glideosome complex and sub-complexes. **(A)** AlphaFold2 model of the composite glideosome complex comprising GAPM1, GAPM2, GAPM3, GAP40, GAP45, GAP50, MyoA, MTIP, and ELC. **(B)** AlphaFold2 model of the GAPM1:GAPM2:GAPM3 heterotrimer in complex with GAP50. **(C)** AlphaFold2 model of the GAP50:GAP45:GAP40 subcomplex. **(D)** AlphaFold2 model of GAP40, GAP45, MyoA, MTIP, and ELC complex. For all models, the Predicted Aligned Error (PAE) colour scale (blue to red) indicates the expected positional error (Å), with lower values (blue) reflecting higher confidence in the relative positioning of residues. Colour-coded bars along the x- and y-axes of the PAE plots denote the boundaries of individual proteins within each model.

**Supplementary Table 3.** Cryo-EM data collection and refinement statistics for GAPM heterotrimer.

|  | GAPMs DDM<br>(EMDB-56647)<br>(PDB 28NA) | GAPMs GDN<br>(EMDB-56677) |
| --- | --- | --- |
| **Data collection and processing** |  |  |
| Magnification | 105,000 | 105,000 |
| Voltage (kV) | 300 | 300 |
| Electron exposure (e<sup>-</sup>/Å<sup>2</sup>) | 39.5 | 40.5 |
| Defocus range (μm) | -1 to -2.4 | -1 to -2.2 |
| Pixel size (Å) | 0.83 | 0.83 |
| Symmetry imposed | C1 | C1 |
| Initial particle images (no.) | 10704382 | 5505208 |
| Final particle images (no.) | 98384 | 181930 |
| Map resolution (Å) | 3.4 | 3.6 |
| FSC threshold | 0.143 | 0.143 |
| Map resolution range (Å) | 2.93-5.50 | 3.02-4.83 |
| **Refinement** |  |  |
| Map sharpening *B* factor (Å<sup>2</sup>) | -46.6217 | -94.2536 |
| Model composition |  |  |
| Non-hydrogen atoms | 5502 |  |
| Protein residues | 683 |  |
| Ligands | 0 |  |
| *B* factors (Å<sup>2</sup>) |  |  |
| Protein | 54.12 |  |
| Ligand | -- |  |
| R.m.s. deviations |  |  |
| Bond lengths (Å) | 0.004 (0) |  |
| Bond angles (°) | 0.546 (0) |  |
| Validation |  |  |
| MolProbity score | 1.38 |  |
| Clashscore | 6.52 |  |
| Poor rotamers (%) | 0.87 |  |
| Ramachandran plot |  |  |
| Favored (%) | 97.93 |  |
| Allowed (%) | 2.07 |  |
| Disallowed (%) | 0.00 |  |
| Model vs. Data |  |  |
| CC (mask) | 0.87 |  |
| CC (box) | 0.66 |  |
| CC (peaks) | 0.58 |  |
| CC (volume) | 0.86 |  |
| Mean CC for ligands | -- |  |

\*CC(masked) between GAPM DDM model and GDN volume map is 0.78

## Supplementary information titles

**Supplementary Video 1:** *Plasmodium berghei* exflagellated male gamete, with GAPM2-GFP (green) and DNA (Hoechst, blue) labelled.

**Supplementary Data 2:** *Pb*GAPM2 and *Pk*GAPM2 mass spectrometry annotated results.

**Supplementary Table 3:** Cryo-EM data collection and refinement statistics for *Pf*GAPM heterotrimer.

**Supplementary Data 4:** Annotated sequence alignment for Extended Data Figure 7

**Supplementary Data 5:** Annotated sequence alignment for GAPM1

**Supplementary Data 6:** Annotated sequence alignment for GAPM2

**Supplementary Data 7:** Annotated sequence alignment for GAPM3

**Supplementary Table 8:** Primers used in this study

