## Supplementary Table 3 for "GAPMs form a heterotrimeric complex bridging the gliding machinery and the cytoskeleton across Plasmodium species"

**Cryo-EM data collection, refinement and validation statistics**

|  | GAPMs DDM  (EMDB-56647)  (PDB 28NA) | GAPMs GDN  (EMDB-56677) |
| --- | --- | --- |
| **Data collection and processing** |  |  |
| Magnification | 105,000 | 105,000 |
| Voltage (kV) | 300 | 300 |
| Electron exposure (e–/Å^2^) | 39.5 | 40.5 |
| Defocus range (μm) | -1 to -2.4 | -1 to -2.2 |
| Pixel size (Å) | 0.83 | 0.83 |
| Symmetry imposed | C1 | C1 |
| Initial particle images (no.) | 10704382 | 5505208 |
| Final particle images (no.) | 98384 | 181930 |
| Map resolution (Å)  FSC threshold | 3.4  0.143 | 3.6  0.143 |
| Map resolution range (Å) | 2.93-5.50 | 3.02-4.83 |
| **Refinement** |  |  |
| Map sharpening *B* factor (Å^2^) | -46.6217 | -94.2536 |
| Model composition  Non-hydrogen atoms  Protein residues  Ligands | 5502  683  0 |  |
| *B* factors (Å^2^)  Protein  Ligand | 54.12  -- |  |
| R.m.s. deviations  Bond lengths (Å)  Bond angles (°) | 0.004 (0)  0.546 (0) |  |
| Validation  MolProbity score  Clashscore  Poor rotamers (%) | 1.38  6.52  0.87 |  |
| Ramachandran plot  Favored (%)  Allowed (%)  Disallowed (%) | 97.93  2.07  0.00 |  |
| Model vs. Data  CC (mask)  CC (box)  CC (peaks)  CC (volume)  Mean CC for ligands | 0.87  0.66  0.58  0.86  -- |  |
